# Bias-mitigated microbiome inference refines coronary artery disease signature

**DOI:** 10.64898/2026.06.04.730260

**Authors:** Luke Honeybrook

## Abstract

Roughly half the cells in the human body are microbial, and changes in these communities are increasingly implicated in cardiovascular, metabolic, and oncological diseases. Yet identifying which taxa truly differ in abundance, differential abundance (DA), is distorted by four major sources of bias: loss of total microbial load, taxa measurement efficiencies, arbitrary pseudocounts required to handle pervasive zeros, and contamination which has recently driven retractions. No existing DA method accounts for all four. Here we introduce BootDA, a non-parametric bootstrap-based method that explicitly models each bias source without data transformations, pseudocounts, parametric assumptions, or assuming that most taxa are non-DA. In semi-parametric simulations preserving the sparsity (>70% zeros) and correlation structure of real 16S amplicon data, BootDA achieved the highest sensitivity among tested methods, including ANCOM-BC2, LinDA, MaAsLin 3, and Wilcoxon tests, while controlling the false discovery rate. Performance was retained in low biomass settings when contamination contributed ∼50% of counts, and without negative controls, indicating *de novo* decontamination capability. Applied to a coronary artery disease cohort, BootDA refined the original signature to two co-enriched genera, *Klebsiella* and *Gemmiger*, and excluded likely contaminants. BootDA is available as an R package and could generalise to other sparse, high dimensional biological data.

## Introduction

By cell count, the human body is approximately 50% non-human cells [1]. This discrepancy is due to the presence of microorganisms, which are widely thought to be essential for homeostasis, occupying for example the intestines, skin, and saliva [1, 2]. However, microbial communities also contribute to human disease, being implicated in for example colon cancer [3], pathological calcification [4], and cardiovascular diseases [5-9], with many therapeutic interventions directed at altering microbiome compositions under investigation [3]. Critical to understanding these disease mechanisms is the ability to detect microbial taxa that differ in abundance between conditions or groups, that is, differential abundance. In microbiome research, such differences are commonly measured using sequencing methods, particularly via the 16S rRNA gene which is present in nearly all bacteria and provides a rapid and cost effective alternative to culturing-based assays and direct cell counting [2]. Yet these approaches suffer a range of experimental and analytical biases that can substantially distort biological conclusions.

Data from these techniques are typically tables of integer counts corresponding to the number of positive identifications of genetic material of taxa—typically >70% of counts are zero [10, 11]. As count tables do not represent raw cell numbers, compositional rather than count-based analyses were recommended. These in turn required the use of log-ratios and related techniques, which often necessitate adding ‘pseudocounts’ to handle zero values [10]. However, the choice of pseudocount is effectively arbitrary, and different choices can substantially alter inferred DA taxa, biasing outcomes [11, 12]. This represents a primary source of statistical modelling bias, though substantial biases also arise from the experimental protocol itself.

As noted above, compositional analyses were used on the assumption that the multiplicative constant linking sequence counts to raw cell counts was lost. However, it was later shown that count tables can be conceptually interpreted as draws from reference frames of differing sizes [13], analogous to using counting squares of different sizes in ecology, and that these differences in reference frame size can be accounted for, up to a scaling factor [13]. Once estimated, this scaling factor, termed sample bias, enables hypothesis testing and fold-change estimation of abundance on a scale proportional to raw cell counts [12-14]. Recovering this value is critical, as species appearing non-DA under compositional inference can differ substantially on this proportional raw scale, where true disease-associated differences and potential therapeutic targets are revealed [15, 16].

Sequencing protocols involve multiple steps, including cell extraction, lysis, and DNA recovery, each of which can introduce taxon-specific efficiencies. Gram-positive bacteria, for example, often have stronger cell walls, reducing lysis efficiency and DNA yield per cell [17]. The efficiencies of each step combine into an overall taxon-specific efficiency bias that acts on all observations of a taxon across samples [11-13, 17]. Because this overall efficiency bias may vary by orders of magnitude between taxa, sequenced counts may misrepresent true abundances to the same extent [17]. Consequently, a taxon appearing highly abundant may in fact be far less abundant, or vice versa, simply because it is measured more or less efficiently, with substantial impacts on the validity of conclusions drawn [17].

Finally, contamination is increasingly recognised as a major bias source, with the potential to skew inference and invalidate biological conclusions [2, 17]. This issue is particularly prominent in low biomass environments, where substantial fractions of counts may derive from, for example, bacteria inadvertently introduced during sample handling. Indeed, entire microbiomes reported to occur in organs, such as the placenta, have since been revised as deriving entirely from contamination [2]. In other cases, decontamination tools have suggested contamination severe enough to reject 50% of all counts in a count table [2]. Contamination counts have been previously modelled as an additive term applied to each taxon across samples, scaled by a sample bias factor [18].

To our knowledge, no inference technique explicitly accounts for all four sources of bias (Table 1). Here we introduce BootDA, a non-parametric method for inferring DA that models multiplicative sample and taxon biases together with additive contamination, without transformations or pseudocounts. We benchmark BootDA against frontier DA methods in semi-parametric simulations that preserve the sparsity and correlation structure of real count tables, including low biomass settings in which contamination contributes ∼50% of counts. BootDA achieves high sensitivity and controls FDR across scenarios, including when total microbial load is unmeasured and negative controls are unavailable, indicating *de novo* decontamination capability. We then apply BootDA to a coronary artery disease (CAD) cohort, refining the reported microbiome signature to two co-enriched genera, *Klebsiella* and *Gemmiger*, while excluding likely contaminants. Together, these results illustrate BootDA’s potential for microbiome inference and, more broadly, for other sparse, high dimensional biological datasets subject to similar biases.

**Table 1.** Bias sources addressed by BootDA versus frontier microbiome DA methods. ✓ = bias explicitly modelled, blank = not modelled. ^1^Bayesian DA methods, such as ALDEx3, typically assume a uniform Dirichlet prior distribution, but the choice is arbitrary and can bias DA outcomes [19], ^2^considered via a built-in pseudocount sensitivity analysis, ^3^valid only when most taxa are non-DA or spike-in taxa with known abundances are provided [11, 20].

|  | Pseudocount bias | Sample bias | Taxa bias | Contamination bias |
| --- | --- | --- | --- | --- |
| BootDA | ✓ | ✓ | ✓ | ✓ |
| ALDEx3 [21, 22] | <sup>1</sup> | ✓ |  |  |
| ANCOM-BC2 [12] | <sup>2</sup> | ✓ | ✓ |  |
| LinDA [14] |  | ✓ |  |  |
| LOCOM [11] | ✓ | <sup>3</sup> | ✓ |  |
| MaAsLin 3 [20] | ✓ | <sup>3</sup> |  |  |

### A model accounting for biases

Regression models are among the most widely used inference methods in biomedical research, due to their ability to handle categorical and continuous variables and provide interpretable effect estimates [23]. Inference is typically simpler if the regression model has a simple additive form [23-25]. However, parametric inference for such forms typically assumes measured variables exhibit constant variance [23-25]. Count table data typically does not and instead exhibits heteroskedascity [26, 27]. It is primarily because of heteroskadescity, or a requirement to stabilise variance [24, 27], that most DA regression methods posit non-additive models that are subsequently transformed to an additive form. For example, LOCOM applies a logistic transform to infer changes in taxa odds ratios [11], and ANCOM-BC2 applies a log transform to infer changes in log-fold abundances [12]. While effective for stabilising variance, such transformations typically require pseudocounts.

To our knowledge, despite some DA methods making use of non-parametric hypothesis testing methods after transformation [11], none have suggested that non-parametric inference could negate the need to apply a transformation entirely, circumventing the introduction of pseudocounts. To this end we posit the following model for count table data (Eqn. 1).

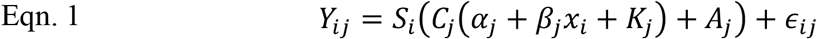

Here, *Y*_*ij*_ is the raw count table reads, *S*_*i*_ is sample bias, *C*_*j*_ is taxa bias, *α*_*j*_ is baseline scaled raw cell counts, *β*_*j*_ is the effect size on scaled raw cell counts due to sample covariate *x*_*i*_, *K*_*j*_ are additional counts due to contamination introduced during experimental stages affected by taxa bias, and *A*_*j*_ are additional counts due to contamination introduced during experimental stages not necessarily affected by taxa bias, and *ϵ*_*ij*_ is non-parametric random noise. Through fitting to multiple resampled datasets, we determine estimates of the effect sizes and their bootstrapped confidence intervals for testing the null hypothesis that *β*_*j*_ ≠ 0, see Methods for full details.

We drew inspiration from ANCOM-BC2 in positing our model (Eqn. 1) and indeed if the contamination terms are discarded it closely resembles the underlying model posited by ANCOM-BC2 [12]. Our model includes multiplicative sample bias and taxa bias terms but, crucially, also includes additive contamination terms that are not necessarily captured by a multiplicative taxa bias term [17]. We follow popular microbiome count table decontamination tools in modelling contamination as a profile across taxa, like a negative control [2, 28]. However, a negative control’s counts cannot be simply subtracted from the count table because it may have a substantially different total microbial load, and hence strong sample bias, which requires accommodating for [2, 28]. Our model accounts for this by incorporating the contamination counts and enabling them to be multiplied by taxa and sample bias factors.

### Comparison to frontier differential abundance methods

Benchmarking DA methods is vital to give confidence for their application in real experiments. To this end, many works have relied on generating synthetic count tables via parametric models, such as negative binomial distributions, before applying spiked-in differences between groups, and recording how many times these were detected [11, 12, 14, 20-22].

However, recent works have noted that parametric generators may not reproduce complex correlations and dependencies observed in real data leading to oversimplified testing scenarios [29-31]. Bootstrap resampling of real datasets to generate synthetic sets, or statistical plasmodes, typically preserves the complexities of the underlying dataset better than parametric generators [29-31]. Indeed, recent work showed several parametric generators produced data that markedly differed in sparsity and variance structure compared to real datasets, and were readily distinguished by machine learning classifiers whereas datasets produced via non-parametric methods were not [30].

The rationale of such non-parametric count table generation may be understood as follows. Taxa counts define an empirical distribution function, and as additional observations are made, new draws from the true underlying distribution are collected, and the empirical distribution function converges to the true distribution [32, 33]. Generating random samples with replacement from this empirical distribution therefore approximates an independent draw from the true distribution [34-36]. This procedure, known as bootstrapping or resampling with replacement, thus yields an approximate independent draw from the true data generating process.

To illustrate the difference, we generated synthetic tables using the popular parametric SparseDOSSA2 and Poisson lognormal mixture model (PLNM) generators [11, 12, 20, 37, 38], which showed marked departures from the original count table (Fig. 1) (see Methods). The PLNM model produced substantially reduced sparsity and a far narrower zero count distribution (Fig. 1A, C), while SparseDOSSA2 more closely matched overall sparsity but showed reduced zero count dispersion (Fi. 1A-B). Both parametric generators also produced visibly different principal component structures compared to the original table (Fig. 1E-F), in agreement with previous work [30]. In contrast, the bootstrap generated table closely reproduced overall sparsity (Fig. 1A, D), the distribution of zero counts (Fig. 1A, D), and the principal component structure (Fig. 1G), indicating better preservation of the empirical distribution and correlations between taxa in the original data.

**Figure 1.**
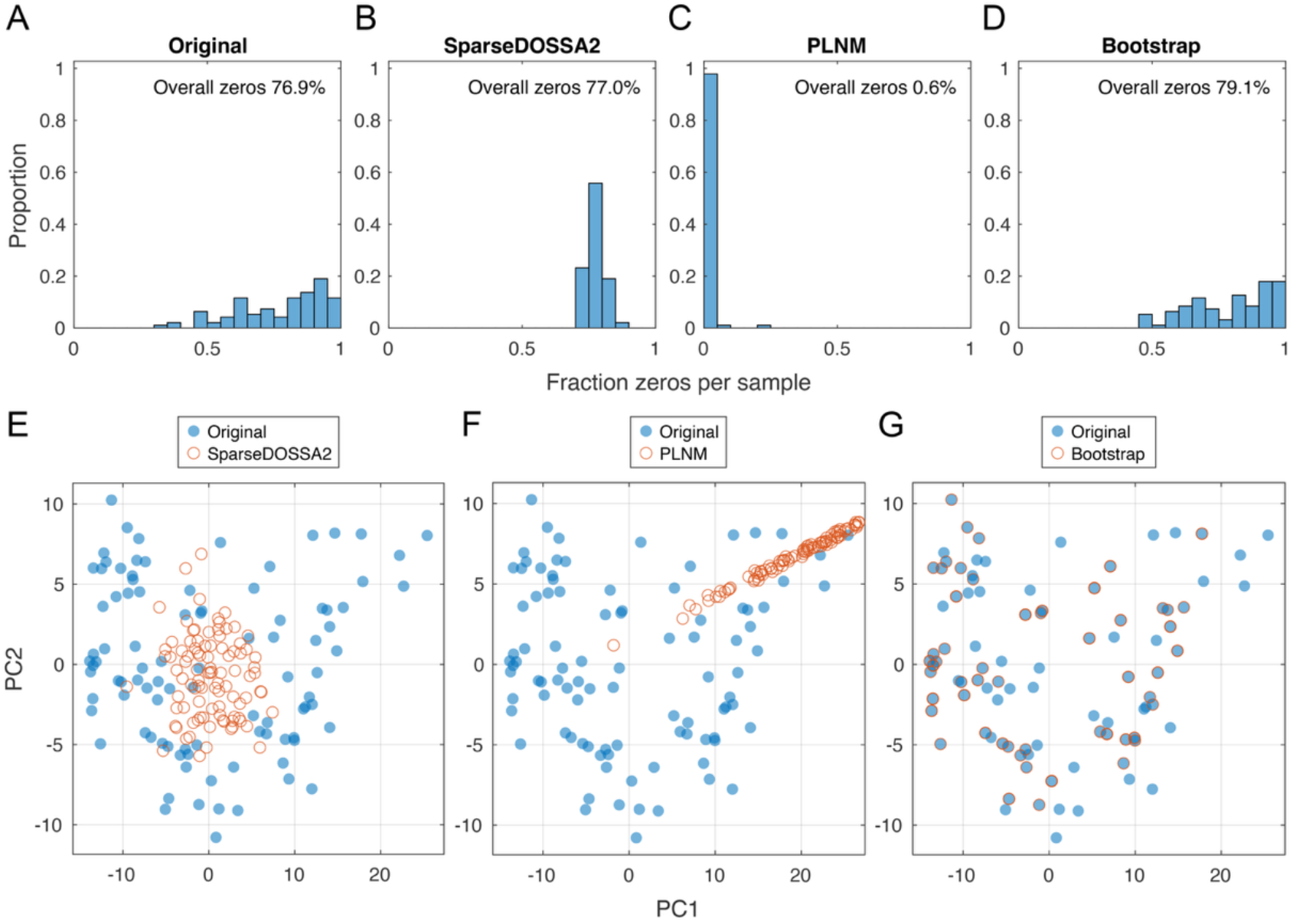
Comparison of synthetic count table generators. Frequency histograms of the fraction of zeros in each sample (row) of (A) the original experimentally derived count table, (B) count table generated by SparseDOSSA2, (C) count table generated by the PLNM model, (D) count table generated by the bootstrap method. Comparison of principal component analyses between the original experimentally derived count table and (E) SparseDOSSA2, (F) the PLNM model, (G) the bootstrap method.

Bootstrapping therefore generates synthetic count tables which can serve as approximately independent experimentally derived datasets. We therefore used this to create a semi-parametric benchmarking protocol whereby baseline count tables were generated via non-parametric bootstrapping followed by spiked-in features. Baseline counts were derived from a gut microbiome dataset from the quantitative microbiome project (QMP) and an upper respiratory tract (URT) dataset, which are popular benchmarking datasets [11, 12, 14, 21, 22], producing baseline count tables with ∼80% and ∼70% zero counts, respectively. We simulated binary, continuous, and multigroup DA settings across a range of sample sizes and spike-in fractions. Similar to previous benchmarking [12], spike-in fractions of 0.1, 0.5, and 0.9 were used to investigate how performance changes when assumptions of most taxa being non-DA, which some methods explicitly assume [11, 12, 20], are challenged.

All synthetic tables additionally incorporated sample and taxa biases and contamination, including a low biomass setting in which contamination contributed approximately 50% of counts. As in previous works [18, 28], contamination was simulated to disproportionately affect prevalent taxa [18, 28]. We further tested scenarios in which sample bias estimates or negative controls were supplied to applicable DA methods. For computational brevity, the full covariate design was assessed only for QMP based simulations, with the remaining scenarios restricted to binary group comparisons. Overall, 11,000 synthetic count tables were generated and analysed per DA method using recommended default settings.

Following previous benchmarking work, we assessed performance via sensitivity, the proportion of truly DA taxa correctly identified, and false discovery rate (FDR), the proportion of detected taxa that were false positives. These are the most widely used performance metrics in DA benchmarking [11, 12, 14, 20-22] and are especially relevant because biological interpretation and identifying therapeutic targets often depends on taxa identified as DA.

We compared BootDA’s performance to several frontier DA methods: ALDEx3 [21, 22], ANCOM-BC2 [12], LinDA [14], LOCOM [11], and MaAsLin 3 [20]. ALDEx3 models count uncertainty through Monte Carlo samples from a Bayesian Dirichlet posterior and sample bias scale uncertainty through a separate model, applies a log-ratio transformation to the combined estimates, and summarises inference across posterior realisations. ANCOM-BC2 forms a log-linear model of observed counts a function of true abundance, sample bias, and taxa bias, aiming to infer covariate effects on absolute abundance after correcting compositional bias. It also supports a pseudocount sensitivity analysis to identify results that depend on the choice of added constant.

LinDA models counts with a linear model after applying a centred log-ratio transform to infer and correct sample bias using the mode of taxa regression coefficients. LOCOM uses logistic regression on taxon pairs, forming and non-parametric permutation testing odds ratios that negate taxa bias and avoid pseudocount use. MaAsLin 3 models prevalence and nonzero abundance separately in a hurdle-style model, enabling separate inference of presence/absence and abundance associations. We also compared against Wilcoxon rank-sum testing on relative abundances, or total sum scaled (TSS) abundances, which is still often used despite its simplicity [5-8, 10, 19, 30]. See Methods for full details.

We found that BootDA generally had the highest sensitivity across QMP covariate scenarios and spike-in fractions, with sensitivity increasing steadily with sample size while maintaining a satisfactorily low FDR (Fig. 2, S1-2). ANCOM-BC2 standard typically achieved a higher sensitivity, but its FDR varied markedly across scenarios and reached as high as 60%. LOCOM performed well at smaller sample sizes, with high sensitivity and low FDR, but its sensitivity plateaued as sample size increased. LinDA was the closest overall competitor to BootDA, showing steadily improving sensitivity and similarly low FDR, although it typically remained ∼30% less sensitive in group comparison settings.

**Figure 2.**
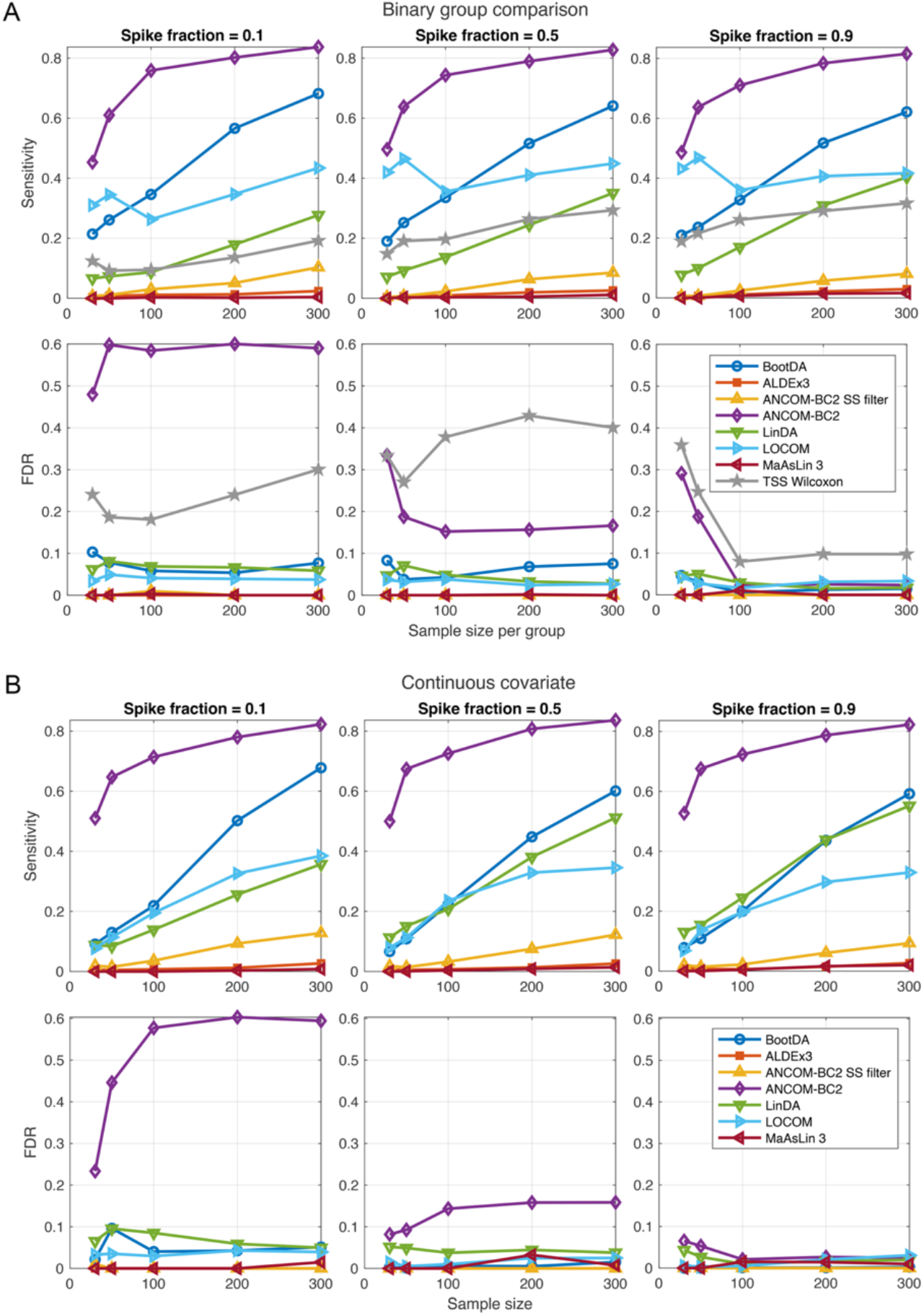
Benchmarking DA methods in simulated (A) binary group and (B) continuous covariate comparisons. Mean sensitivity FDR as a function of sample size per group, for BootDA, ALDEx3, ANCOM-BC2 (standard and pseudocount sensitivity filtered), LinDA, LOCOM, and MaAsLin 3. Wilcoxon rank-sum testing on relative abundances (TSS Wilcoxon) is also performed for the binary group comparison. Means are taken from results over 100 semi-parametric generated count tables derived from the QMP dataset, for spike-in fractions of 0.1, 0.5, 0.9 of taxa.

By contrast, ALDEx3, ANCOM-BC2 with pseudocount sensitivity filter, and MaAsLin3 all maintained extremely low FDRs, generally below 1%, but at the cost of very low sensitivity, typically below 20%. TSS Wilcoxon showed intermediate sensitivity, but like standard ANCOM-BC2 exhibited highly unstable FDR, reaching up to ∼40%. Similar overall patterns were observed in the URT simulations, although sensitivities were generally higher across methods while FDR trends were largely unchanged (Fig. 2A, S3). Notably, the elevated and unstable FDR of ANCOM-BC2 and Wilcoxon persisted and did not clearly improve with increasing sample size (Fig. S3).

We next evaluated scenarios in which total microbial load was available, for example from flow cytometry or machine learning prediction [15, 16], allowing known sample bias to be supplied to the model. Among comparator DA methods, only ALDEx3 and MaAsLin3 could incorporate prior sample bias information, and we additionally considered Wilcoxon rank-sum tests after rescaling counts by the known sample biases.

Providing sample bias information substantially altered performance relative to the setting in which microbial load was unavailable (Fig. 3A). For BootDA, sensitivity decreased modestly by around 5% across scenarios, with a slight increase in FDR. In contrast, ALDEx3 showed a marked increase in sensitivity when given the prescribed sample biases, even exceeding BootDA at larger sample sizes when the spike-in fraction was 0.9, although its FDR remained unstable and exceeded 10% when the spike-in fraction was 0.5. MaAsLin3 showed little change in sensitivity, but its FDR increased modestly. Rescaling counts by known sample bias before applying the Wilcoxon rank-sum test reduced both sensitivity and FDR substantially, in some cases by as much as 20%.

**Figure 3.**
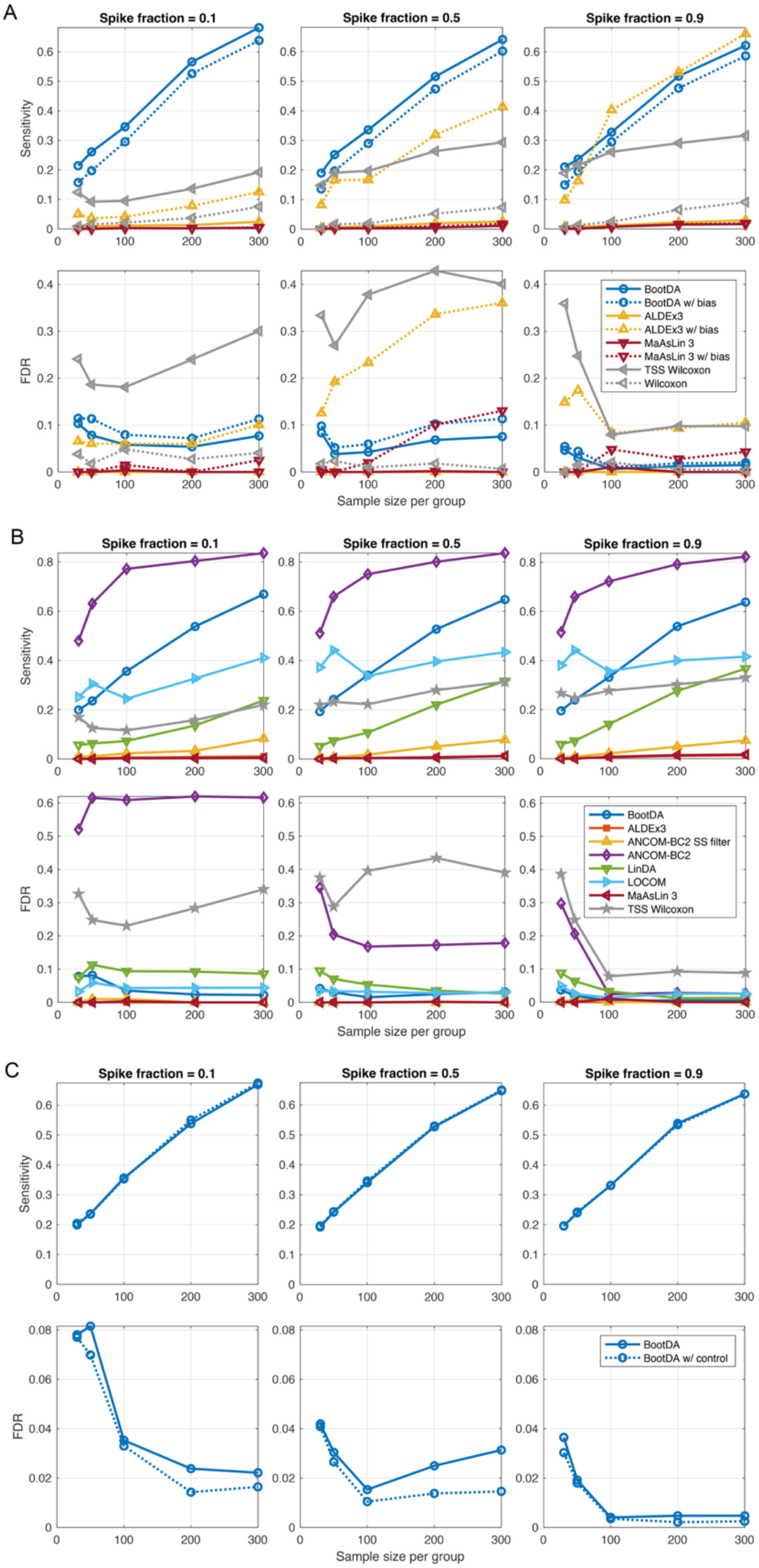
Performance of DA methods under additional bias conditions. Mean sensitivity and FDR as a function of sample size per group for BootDA and comparator methods across three settings: (A) simulations in which true sample biases were supplied (w/ bias) or withheld from applicable methods, (B) a low biomass scenario in which contamination accounted for approximately 50% of all counts, and (C) simulations in which a corresponding negative control was supplied (w/ control) or withheld from BootDA in the low biomass setting.

In the low biomass scenario, method sensitivities and FDRs were broadly similar to those in the original setting (Fig. 2A, 3B), with BootDA again generally achieving higher sensitivity than competing methods, except at smaller sample sizes, while maintaining FDR control. We also evaluated scenarios in which a corresponding negative control was available for each count table of the low biomass setting. Comparator DA methods were not evaluated in this setting because none explicitly model contamination. In this scenario, BootDA’s sensitivity was essentially unchanged, but its FDR decreased slightly (Fig. 3C).

In summary, across nearly all simulated scenarios BootDA showed the highest sensitivity of any comparator method whilst maintaining FDR control. Notably, this advantage persisted even in contaminated settings, including the low biomass scenario in which approximately 50% of all counts were attributable to contamination. Motivated by this robust performance, we next applied BootDA to a CAD gut microbiome dataset.

Means are taken from results over 100 semi-parametric generated count tables derived from the QMP dataset, for spike-in fractions of 0.1, 0.5, 0.9 of taxa.

### Application to coronary artery disease

We applied BootDA to a gut microbiome dataset from Liu et al. comprising faecal 16S rRNA gene amplicon profiles of the V3-V4 region from a healthy cohort (n = 52) and those from pooled CAD cases (n = 186) spanning stable CAD (n = 55), unstable angina (n = 91), and myocardial infarction (n = 40) [5]. These classifications of CAD are all manifestations of atherosclerosis, a process of calcified plaque formation that restricts blood flow [9].

BootDA returns a scaled effect size, *C*_*j*_*β*_*j*_, equal to the taxa bias multiplied by the effect size on scaled raw cell counts per unit change in the covariate (see Methods). For a categorical covariate, this corresponds to the difference between groups. BootDA identified three CAD-enriched taxa (Fig. 4A): the phylum Firmicutes (scaled effect size = 45.9, 95% Bonferroni-Dunn adjusted bootstrap percentile CI [1.0, 135.4]), the genus *Klebsiella* (scaled effect size = 312.4, [1.4, 929.5]) and the genus *Gemmiger* (scaled effect size = 261.7, [38.3, 587]) as enriched in CAD relative to healthy controls.

**Figure 4.**
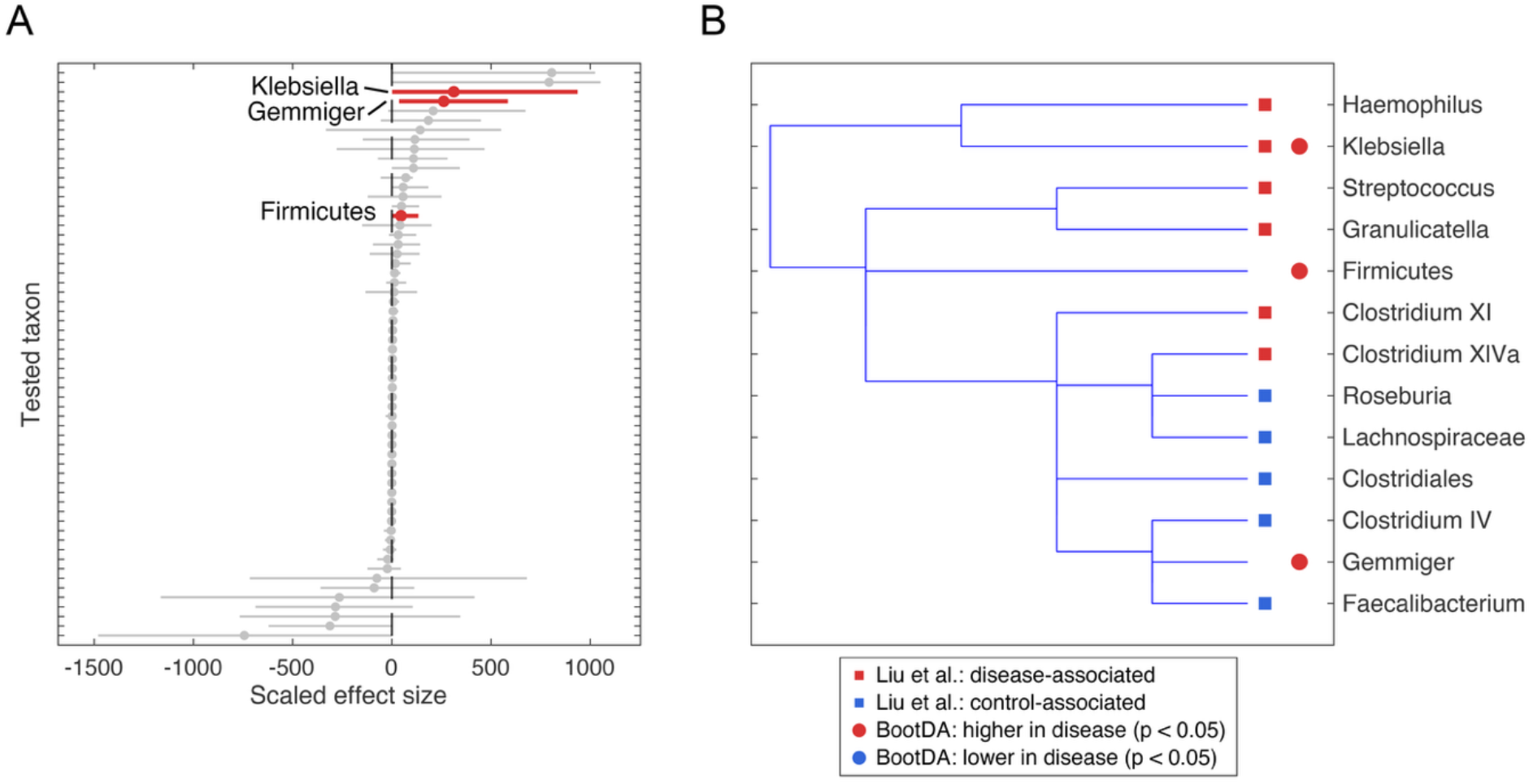
BootDA identifies a refined set of CAD-associated taxa in the Liu et al. [5] dataset. A) Scaled effect sizes (*C*_*j*_*β*_*j*_) with 95% Bonferroni-Dunn adjusted bootstrap percentile CIs estimated by BootDA for all tested taxa between healthy controls (n = 52) and pooled CAD cases (n = 186). Taxa reaching statistical significance (p < 0.05) are highlighted in red (CAD-enriched). B) Comparative phylogenetic dendrogram illustrating taxa identified as CAD- or control-associated by the original Liu et al. [5] analysis and by BootDA. Filled squares denote associations identified in the original analysis (red: CAD-associated, blue: control-associated), filled circles denote BootDA significant taxa (red: higher in CAD, p < 0.05, blue: lower in CAD, p < 0.05).

Compared to the original analysis, which identified a larger set of taxa as healthy and CAD-associated, BootDA identified a much smaller number of CAD-associated taxa (Fig. 4B). This is consistent with the prevailing hypothesis that only a small number of taxa are truly DA in most diseases [11-14]. The identification of *Klebsiella* as CAD-enriched agreed with Liu et al.’s original analysis (Fig. 4B). The broad Firmicutes feature was less specific and *Gemmiger* was not identified previously [5]. Also of note, BootDA did not identify taxa typically associated with kit contamination such as *Streptococcus* as DA, whereas Liu et al.’s original analysis did [5].

## Discussion

We developed a method, BootDA, for microbiome sequencing counts that, unlike other frontier DA methods, accounted for pseudocount bias, sample bias, taxa efficiency bias, and contamination counts simultaneously.

We found that BootDA generally showed the strongest combination of sensitivity and FDR control against comparator methods across realistic semi-parametric simulations. We found that other frontier DA methods typically controlled FDR but at the cost of lower sensitivity or attained higher sensitivity only with markedly unstable FDR, a trade-off that is difficult to justify when mechanistic biological interpretations depend on the credibility of the taxa identified as DA [29, 30]. Of particular note was that Wilcoxon rank-sum tests performed on relative counts (TSS Wilcoxon), despite its simplicity and continued use in the literature [5, 10, 30], showed markedly unstable and high FDRs. Previous works have also shown this [11, 13, 14].

An important point of our semi-parametric benchmark protocol is that it was challenging and realistic. It made use of non-parametric baseline count table generators that preserve sparsity and complex correlations of real data, and then included realistic sample, taxa, and contamination biases, forming a statistical plasmode [31]. Indeed, to our knowledge this is the first study to explicitly include and examine how all these sources of biological and experimental bias simultaneously challenge DA methods. Recent benchmarking work has argued that plasmode or bootstrap-based strategies preserve empirical sparsity, dependence structure, and other properties of real microbiome data more faithfully than fitted parametric generators that have been used to assess DA methods previously [29-31]. Our comparison with PLNM and SparseDOSSA2 was consistent with that view, with bootstrap generated tables more closely reproducing zero count structure and principal component geometry than parametric alternatives. Hence, sensitivity and FDR performance observed here likely better reflect the behaviour of DA methods under experimentally plausible conditions than benchmarks based on parametric generators.

The difference between the QMP and URT simulations also helps clarify where BootDA’s advantage appears strongest. Sensitivities were generally higher in the URT setting across methods, in particular parametric methods, which is likely due to the lower sparsity of the URT baseline table relative to QMP (∼70% vs ∼80%). Fewer zeros reduce heteroskedasticity and stabilise variance, on which parametric models depend more heavily for inference than non-parametric methods [14, 26, 27]. This observation suggests that BootDA’s strongest performance over other methods lies in very high sparsity datasets common in amplicon-based microbiome studies.

Where sample bias was supplied to DA methods able to use it, inference did not substantially improve. For BootDA, providing true sample biases had a negligible effect on sensitivity and slightly elevated FDR, whilst in other techniques the same information degraded and destabilised FDR control. This suggests that normalisation by total microbial load does not necessarily lead to improved inference in the presence of other confounding effects such as taxa efficiency bias and contamination. The result is practically useful because previous work has proposed quantifying total microbial load to provide accurate DA inference [15, 16]. Our work suggests that such measurements may not be necessary if a method explicitly accounts for other biases.

In the low biomass scenario, where contamination accounted for approximately 50% of counts, BootDA achieved similar or higher sensitivities to other methods whilst maintaining FDR control. This is crucial because contamination is now widely recognised as a primary problem in microbiome inference, particularly in low biomass environments, yet to our knowledge no DA method explicitly models contamination, and existing decontamination tools either identify taxa with contaminants but cannot correct the count table [2, 28] or estimate and subtract contaminant counts without considering other biases [18]. Previous work has suggested that prevalence-based methods could mitigate the difficulties of contamination counts [11, 20], but our results show that this is not sufficient on its own. Indeed, MaAsLin 3, which incorporates both abundance and prevalence modelling explicitly [20], did not approach BootDA’s sensitivity in our simulations. Equally notable, supplying a negative control to BootDA only marginally improved its performance over the case when a negative control was not supplied, indicating a de novo contamination identification and mitigation ability, similar to some other tools [28]. This could be very useful as negative controls are often unavailable in historical and public datasets where BootDA may be readily applied [2, 28].

A further practical advantage of BootDA is that it requires no prevalence filter, that is, no preprocessing step that removes taxa present in only a small fraction of samples. Many existing DA methods use or recommend such filters on the assumption that low prevalence taxa may be contamination artefacts [12]. However, rare taxa may hold important information as abundance below detection thresholds does not preclude a taxon from having meaningful impacts on health, as some microbial species can cause infection even at very low doses [20]. Indeed, MaAsLin 3 explicitly recommends against prevalence filters for these reasons [20]. Further, the choice of prevalence threshold is somewhat arbitrary, with values typically ranging from 10-20% [11, 12]. That BootDA does not require a prevalence filter is therefore beneficial not only because no potentially informative taxa are discarded, but also because it mitigates prevalence filter threshold bias. Also, unlike approaches that present only a p-value, BootDA produces scaled effect sizes and confidence intervals that help illustrate abundance changes and estimation uncertainty, in line with recommendations for more comprehensive statistical reporting [12, 13, 39].

We applied BootDA to a 16S rRNA gene sequencing dataset of healthy and CAD cohorts that lacked negative control [5]. Liu et al.’s original analysis reported numerous DA taxa using Wilcoxon rank-sum testing of relative abundances [5]. Reanalysis with BootDA identified three CAD-enriched taxa (all p < 0.05): the phylum Firmicutes and the genera *Klebsiella* and *Gemmiger*. Because *Gemmiger* was the only Firmicutes genus identified as DA, we attribute the phylum-level enrichment to *Gemmiger* enrichment, leaving two genus-level DA identifications. This is consistent with the hypothesis that only a small number of taxa are truly DA in most diseases [11-14]. Notably, *Streptococcus* and other genera found to be CAD-enriched by Liu et al. are also commonly reported kit and reagent contaminants [2, 28], and were not identified by BootDA.

To our knowledge, the application of bias-mitigating DA methods to cardiovascular-gut microbiome inference is rare. Previous works on DA in CAD have typically relied on Wilcoxon tests on relative abundances and similar methods that do not explicitly account for biases. Of the few works that have used bias-mitigating methods (Table 1), results are conflicting. *Klebsiella* has been suggested as enriched in CAD, though only at the family level [7], as depleted in CAD [8], and as neither enriched nor depleted [6]. No works using bias-mitigating methods have proposed an association with *Gemmiger*. The co-enrichment of *Klebsiella* and *Gemmiger* identified here refines these previous results and suggests a simpler CAD-gut microbiome signature.

The *Klebsiella* genus comprises opportunistic bacteria that can promote inflammation, and may contribute to CAD through release of inflammatory products into the circulation, endothelial activation, and systemic immune activation [9]. *Gemmiger*, by contrast, is a less well characterised anaerobic genus whose enrichment may reflect altered microbial metabolic activity. Their co-enrichment could reflect parallel responses to the complex CAD host environment, such as chronic vascular inflammation and medication exposure. Or alternatively, the two genera may act cooperatively, with *Klebsiella* driving an inflammatory state and *Gemmiger* influencing metabolic processes that affect immune activation, microbial metabolite production, and cholesterol handling, consistent with proposed immune-metabolic mechanisms of gut microbial influence in disease [3, 9]. The association we identify is therefore informative both as a diagnostic signature of CAD-associated host physiology and as a guide for mechanistic investigation, for instance via *in vitro* models of CAD [4].

Overall, BootDA represents a powerful method for identifying truly DA taxa in count data distorted by pseudocount, sample, taxa efficiency, and contamination biases. We found it generally achieved the highest sensitivities whilst controlling FDR across a range of simulated scenarios that challenged other frontier DA methods. Though developed for microbiome count tables, the biases addressed by BootDA are common in sparse, high-dimensional biological datasets suggesting its broader applicability to other omics datasets, such as single-cell transcriptomics [10, 14, 20, 21, 26, 29]. As illustrated by its application to a historic cardiovascular disease dataset, BootDA may clarify and uncover previously undetected disease-associated taxa, supporting more reliable microbiome discovery and accelerating the development of diagnostic and therapeutic strategies. We have implemented our method in the R package BootDA, which is available on GitHub at https://github.com/luke-h-code/BootDA.

## Methods

### Hypothesis testing in microbiome data with bias correction

In Eqn. 1 we posited the original model. Eqn. 1 can be rewritten as Eqn. 2.

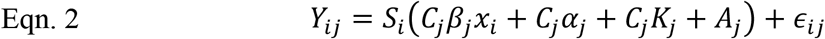

We introduce the terms *m*_*j*_ = *C*_*j*_*β*_*j*_ and *c*_*j*_ = *C*_*j*_*α*_*j*_ + *C*_*j*_*K*_*j*_ + *A*_*j*_. Substitution of these in to Eqn. 2 gives Eqn. 3.

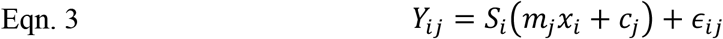

We wish to test the hypothesis that *β*_*j*_ differs with covariate *x*_*i*_. Following previous works [11, 12, 17], we assume taxa bias does not change with covariate. Hence, taxa bias is a row vector *C*_*j*_ and so testing the hypothesis that *β*_*j*_ ≠ 0 is equivalent to testing that *C*_*j*_*β*_*j*_ ≠ 0, and hence equivalent to testing *m*_*j*_ ≠ 0.

Eqn. 3 may be seen as a linear regression in which each row is scaled by an unknown constant, *S*_*i*_. Evaluating the hypothesis *m*_*j*_ ≠ 0 therefore requires estimating both *m*_*j*_ and *S*_*i*_.

We estimate *S*_*i*_ by treating Eqn. 3 as a separable partially linear least squares problem with two parameter blocks: (1) *S*_*i*_ and (2) *m*_*j*_, *c*_*j*_. These are estimated iteratively by alternating least squares (ALS) where one block is held fixed while the other is updated to minimise the residual sum of squares and alternating in this way until convergence [40].

We first write Eqn. 3 in matrix notation (Eqn. 4).

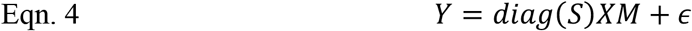

Where *Y* has dimensions n rows (samples) and d columns (taxa), *diag*(*S*) is the diagonalised matrix of the column vector of sample biases with n rows, *M* is the coefficient matrix with d columns 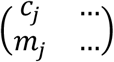, and *X* is the design matrix with n rows like 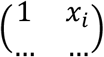. Rows of *Y* in which all counts are zero (i.e. samples with no observed taxa) are removed before fitting, as the ALS update for *S* is undefined for such rows.

To assess the sampling variation of the parameter estimate of interest, 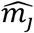, we employed bootstrap resampling. Bootstrap count tables were constructed by sampling rows of *Y* (jointly with *X*) with replacement, with resampling performed within groups when the covariate was categorical so that the observed group structure was preserved, and across all samples when the covariate was continuous [36, 41]. The resulting empirical distribution of the bootstrap estimates of *m*_*j*_ were used to construct their confidence intervals. Because many taxa are tested simultaneously, false positives are expected by chance [23, 29]. We therefore applied the Bonferroni-Dunn correction which adjusts the significance (α) level via *α*_*adj*_ 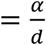[42,43], where we set α as 5% (0.05) and d is the number of taxa being tested. For each taxon, a (1 − *α*_*adj*_) percentile confidence interval for *m*_*j*_ was constructed from the bootstrap distribution. The number of bootstrap samples (B) to form the confidence interval is chosen such that 100 samples are expected in each tail i.e. 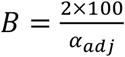. A taxon was declared DA if its confidence interval excluded zero.

The iterative technique we used for each bootstrap sample:

1. Propose initial *Ŝ* as the row-wise mean of *Y*, then mean normalise.
2. Update 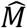 using ordinary least squares on the rescaled response, *diag*(*S*)^−1^*Y*, (i.e. minimise the sum of squared residuals of the rescaled response), which has standard closed form 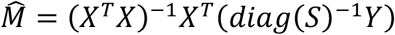 [44].
3. Update 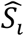 by minimising the sum of squared residuals. With *M* fixed, this is regression of *Y*_*ij*_ through the origin on 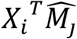 which has the standard closed form 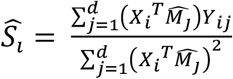 [44]. Then mean normalise.
4. To mitigate misattribution of signal to *S* instead of *M*, posit that log(*Ŝ*) may become linearly related to the covariate design matrix such that log(*Ŝ*) = *Xγ* + *ε*, where *γ* is a temporary coefficient set. The log transform of *Ŝ* does not require pseudocounts because *Ŝ* is always positive by construction. Estimate *γ* by ordinary least squares and subtract the fitted contribution of the non-intercept covariates, *X*_−1_*γ*_−1_, from log(*Ŝ*) and then exponentiate i.e. *Ŝ* = exp(log(*Ŝ*) − *X*_−1_*γ*_−1_). This process is often called residualisation and is similar to how limma removes batch effects [26]. Finally, mean normalise.
5. Repeat steps 2-4 until the relative change in the total sum of squares between successive iterations falls below a prescribed tolerance, or the number of iterations exceeds a prescribed limit. Final 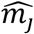 coefficients are recorded.

### Synthetic count table generation

Taxa count tables from the QMP dataset [15] were obtained as in [21]. Raw sequencing data were downloaded from the European Nucleotide Archive (accession PRJEB21504) and processed into a sequence variant table using DADA2 [45] with recommended defaults: filterAndTrim with maxEE = 4, default error-rate learning, core denoising, paired-end merging, and consensus chimera removal. Taxonomy was assigned with the RDP classifier [46]. We only considered the diseased cohort and, following [12], a set of 100 taxa were chosen. This set was taken to be the 100 most prevalent taxa, yielding ∼80% zeros (Fig. 1A). The left oropharyngeal microbiome count table from the URT dataset [47] was obtained from the LOCOM R package [11]. To ensure high sparsity we sampled the most prevalent 250 taxa which yielded ∼70% zeros.

From these original count tables, all subsequent simulations were generated. The derived QMP and URT count tables were bootstrap sampled to create independent count table experiments. Three covariate designs were considered: a binary group design, a continuous covariate design, and a three group design with spike-in taxa DA in groups 2 and 3 only. Group sizes of 30, 50, 100, 200, and 300 samples per group were simulated, with 100 bootstrap replicates generated for each parameter combination.

DA was introduced by applying taxon-specific spike-in fold-changes (*f*_*j*_), sampled with replacement from (−10, −5, −2, 2, 5, 10), to a specified fraction of taxa (0.1, 0.5, or 0.9) similar to [12]. Eligible taxa were required to have at least eight nonzero counts in every relevant group. For categorical designs, spike-ins were applied only within non-reference groups. For continuous designs, a standard Gaussian predictor *x*_*i*_ was generated and effects imposed multiplicatively as *exp*(*a*_*j*_*x*_*i*_), where *a*_*j*_ = *sign*(*f*_*j*_)*log*|*f*_*j*_|, similar to [12]. Taxa bias factors were sampled uniformly from 0.1 to 1.0 and applied across taxa, as in [12]. A contamination profile was then added to each sample by selecting one source sample at random from the QMP or URT table and scaling its counts to 5% of the original values (similar to [28]). In the low biomass simulations, counts were instead scaled to 50%. Finally, sample bias factors, bootstrap-sampled from mean normalised total microbial loads measured by flow cytometry in the QMP diseased cohort [15], were applied multiplicatively to the counts. Final counts were rounded to the nearest integer.

For Fig. 1, features of the derived QMP count table were compared with those of a single bootstrap sample and with single synthetic count tables generated using the PLNM and SparseDOSSA2, each fitted to the derived QMP count table. The PLNM model [38], implemented in the ANCOMBC R package [12], was run without prevalence filtering, with the library size mean set to that of the derived count table. SparseDOSSA2 [37] was fitted once to the observed count table using a regularisation parameter of 1 and a maximum of 100 fitting iterations, after which a synthetic count table was generated from the fitted model with the target median sequencing depth matched to that of the derived QMP count table.

### Differential abundance methods

DA was evaluated using BootDA, as described earlier, ANCOM-BC2, MaAsLin3, ALDEx3, LOCOM, and LinDA. For each simulated dataset and comparison, the count table was subset to the samples relevant to the comparison of interest and supplied to each method using the corresponding binary group indicator or continuous covariate as the sole explanatory variable. Sensitivity and FDR were calculated from taxa at a significance threshold of α = 0.05.

For ANCOM-BC2, analyses were run using the authors’ recommended Holm-Bonferroni multiple comparison adjustment [48], restriction to taxa prevalent in at least 10% of samples, and 5% percentile small positive constant to add to the test statistic [12]. Both the standard and pseudocount sensitivity filtered ANCOM-BC2 methods were used and their results recorded separately. For MaAsLin 3, analyses were run using the authors’ recommended [20] TSS normalisation, Benjamini-Hochberg multiple comparison adjustment, and no restriction on taxa prevalence [49]. Taxa determined to be DA were pooled from both the abundance and prevalence methods.

For ALDEx3, analyses were run using the authors’ recommended [21, 22] linear modelling framework with 1000 Monte Carlo Dirichlet samples, Benjamini-Hochberg multiple comparison adjustment, and the centred log-ratio (clr) scale model, which generalises clr normalisation by allowing uncertainty in the underlying scale assumption, with γ = 0.5. For LOCOM, analyses were run using the authors’ recommended [11] restriction to taxa prevalent in at least 20% of samples, with permutation-based inference performed using up to 1,000 permutations in the group comparison case. For continuous covariates, permutation testing is not recommended. However, when we disabled permutation testing, errors occurred owing to an apparent bug in the LOCOM code. A similar problem was noted in the peer review file of [12]. Hence, we used categorical variable parameters instead. For LinDA, analyses were run using the recommended [14] no restriction on taxa prevalence, Benjamini-Hochberg multiple comparison adjustment, and 97% winsorization.

For total sum scaled Wilcoxon comparison, analyses were run by row normalising each count table and then performing standard two sided Wilcoxon signed-rank tests on each taxa with Benjamini-Hochberg multiple comparison adjustment. This method was not performed for the continuous covariate scenarios.

In the supplied sample bias scenario, BootDA, MaAsLin 3 and ALDEx3 were supplied the prescribed sample biases. In the case of BootDA, these were used as the initial *Ŝ* instead of step 1 of the ALS scheme. In the case of ALDEx3, count tables were log2 transformed as per the authors’ recommendations [21, 22]. For the Wilcoxon rank-sum case, counts were multiplied by the prescribed sample biases and no TSS was applied. In the supplied negative control case, BootDA’s ALS scheme initially begins at step 2 and the initial guess for the *c*_*j*_ values are set to the counts of the negative control.

Across methods, taxa not tested because of method-specific internal filtering were retained in the denominator when computing sensitivity and false discovery rate, but were treated as non-discoveries. Results were calculated separately for each simulated dataset and then summarised as the mean across the 100 replicate datasets generated under each parameter combination.

All benchmarking code used is provided in the Supplementary Code files. BootDA was implemented in MATLAB for the simulation tests, then translated to R (available on GitHub) for application to the CAD dataset, as R is more widely used for microbiome statistical analysis. Both implementations perform identical computations, but because random number generation differs between MATLAB and R, bootstrapped confidence intervals may show small numerical differences.

### Coronary artery disease dataset

Raw paired-end 16S rRNA gene sequencing data for the Liu et al. CAD cohort were obtained from the Sequence Read Archive (BioProject PRJNA503710, accession SRP167862) via ENA. A sequence variant table was generated using DADA2 [45] with the default paired-end workflow: filterAndTrim with maxEE = c(4,4), default error-rate learning and denoising, paired-end merging, and chimera removal via removeBimeraDenovo (consensus method). Taxonomy was assigned with the RDP classifier [46] on the RDP training set, with a minimum bootstrap threshold of 80% to match Liu et al.’s [5]. Species-level assignment via addSpecies recovered no exact matches, so we restricted taxa DA analysis to the genus level and above. The discovery and validation sub-cohorts were combined, and CAD classifications were pooled into a single CAD group, giving 52 healthy and 186 CAD samples for DA testing.

## Author contributions

L.H. conceived the paper, derived the models, coded and performed simulations, and wrote and revised the manuscript. S.B. supervised.

## Acknowledgements

British Heart Foundation grant FS/4yPhD/F/20/34134 (LH). LH acknowledges the use of the UCL Myriad High Performance Computing Facility (Myriad@UCL), and associated support services, in the completion of this work.

## Data Availability Statement

All benchmarking code used is provided in the Supplementary Code files. BootDA is available as a publicly accessible R package on GitHub at https://github.com/luke-h-code/BootDA.

