## Supplementary Code for "Bias-mitigated microbiome inference refines coronary artery disease signature": simulation_eval_common.R

```
#!/usr/bin/env Rscript

suppressPackageStartupMessages(library(tidyverse))

# Force all BiocParallel calls (e.g. inside ANCOMBC2) to run serially,
# matching MATLAB's maxNumCompThreads(1) / pctRunOnAll maxNumCompThreads(1).
# Without this, SnowParam spawns hidden child R processes on Windows even
# when n_cl = 1, multiplying memory use across concurrent Rscript instances.
if (requireNamespace("BiocParallel", quietly = TRUE)) {
  BiocParallel::register(BiocParallel::SerialParam())
}

`%||%` <- function(x, y) if (is.null(x) || length(x) == 0) y else x

find_latest_checkpoint <- function(out_dir, stem) {
  cps <- list.files(out_dir, pattern = paste0("^", stem, "_checkpoint_[0-9]{4}\\.csv$"), full.names = TRUE)
  if (!length(cps)) return(NULL)
  cps[[which.max(suppressWarnings(as.integer(sub(".*_checkpoint_([0-9]{4})\\.csv$", "\\1", basename(cps)))))]]
}

parse_index_list <- function(x) {
  if (is.na(x) || x == "") return(integer(0))
  out <- suppressWarnings(as.integer(strsplit(x, ";", fixed = TRUE)[[1]]))
  out[!is.na(out)]
}

parse_numeric_list <- function(x) {
  if (is.na(x) || x == "") return(numeric(0))
  out <- suppressWarnings(as.numeric(strsplit(x, ";", fixed = TRUE)[[1]]))
  out[!is.na(out)]
}

resolve_dataset_path <- function(path_value, meta_file) {
  if (is.na(path_value) || !nzchar(path_value)) stop("Encountered empty output_csv path in metadata.")
  if (grepl("^/", path_value)) return(path_value)
  candidates <- c(
    file.path(dirname(meta_file), path_value),
    file.path(dirname(dirname(meta_file)), path_value),
    file.path(dirname(meta_file), basename(path_value))
  )
  hit <- candidates[file.exists(candidates)]
  normalizePath((hit %||% candidates)[[1]], winslash = "/", mustWork = FALSE)
}

get_taxa_columns <- function(dat) {
  meta_cols <- c("synthetic_sample_id", "source_sample_id", "group", "continuous_covariate")
  names(dat)[vapply(dat, is.numeric, logical(1)) & !(names(dat) %in% meta_cols)]
}

parse_group_betas <- function(mrow, group_id, n_taxa) {
  idx <- parse_index_list(mrow[[paste0("spike_taxa_indices_group", group_id)]])
  beta <- parse_numeric_list(mrow[[paste0("spike_beta_group", group_id)]])
  n <- min(length(idx), length(beta))
  out <- rep(0, n_taxa)
  if (!n) return(out)
  idx <- idx[seq_len(n)]
  keep <- idx >= 1 & idx <= n_taxa
  out[idx[keep]] <- beta[seq_len(n)][keep]
  out
}

parse_sample_loads <- function(mrow, row_idx) {
  if (!"sample_bias_factors" %in% names(mrow)) stop("Metadata is missing required column: sample_bias_factors")
  loads_all <- parse_numeric_list(mrow$sample_bias_factors[[1]])
  if (!length(loads_all)) stop("Dataset ", mrow$dataset_id[[1]], " has no sample_bias_factors values.")
  if (max(row_idx) > length(loads_all)) stop("Dataset ", mrow$dataset_id[[1]], " sample_bias_factors are too short.")
  loads <- loads_all[row_idx]
  if (any(!is.finite(loads)) || any(loads <= 0)) stop("Dataset ", mrow$dataset_id[[1]], " has invalid supplied total microbial loads.")
  loads
}

build_dataset_comparisons <- function(mrow, dat, taxa_cols, sample_load_mode = FALSE) {
  scenario <- as.character(mrow$scenario_name[[1]])
  n_taxa <- length(taxa_cols)
  beta_by_group <- lapply(1:3, function(g) parse_group_betas(mrow, g, n_taxa))
  add_loads <- function(x, idx) if (sample_load_mode) c(x, list(sample_loads = parse_sample_loads(mrow, idx))) else x
  if (scenario == "continuous_covariate") {
    if (!"continuous_covariate" %in% names(dat)) stop("Dataset ", mrow$dataset_id[[1]], " is missing 'continuous_covariate'.")
    effect_values <- suppressWarnings(as.numeric(dat$continuous_covariate))
    if (anyNA(effect_values)) stop("Dataset ", mrow$dataset_id[[1]], " has non-numeric continuous covariate values.")
    row <- list(
      comparison_name = "continuous_effect",
      reference_group = NA_real_,
      target_group = NA_real_,
      row_idx = seq_len(nrow(dat)),
      effect_values = effect_values,
      truth = abs(beta_by_group[[1]]) > 0
    )
    return(list(add_loads(row, seq_len(nrow(dat)))))
  }
  if (!"group" %in% names(dat)) stop("Dataset ", mrow$dataset_id[[1]], " is missing 'group'.")
  group_values <- suppressWarnings(as.numeric(as.character(dat$group)))
  if (anyNA(group_values)) stop("Dataset ", mrow$dataset_id[[1]], " has non-numeric group labels.")
  pair_list <- switch(
    scenario,
    binary_covariate = matrix(c(1, 2), ncol = 2, byrow = TRUE),
    three_group_spikein = matrix(c(1, 2, 1, 3, 2, 3), ncol = 2, byrow = TRUE),
    stop("Unsupported scenario_name: ", scenario)
  )
  lapply(seq_len(nrow(pair_list)), function(i) {
    ref <- pair_list[i, 1]
    tgt <- pair_list[i, 2]
    keep <- which(group_values %in% c(ref, tgt))
    add_loads(list(
      comparison_name = paste0("group", tgt, "_vs_group", ref),
      reference_group = ref,
      target_group = tgt,
      row_idx = keep,
      effect_values = as.numeric(group_values[keep] == tgt),
      truth = abs(beta_by_group[[tgt]] - beta_by_group[[ref]]) > 0
    ), keep)
  })
}

calc_sens_fdr <- function(truth, calls) {
  if (!is.null(names(truth)) && !is.null(names(calls)) && any(nzchar(names(truth))) && any(nzchar(names(calls)))) calls <- calls[names(truth)]
  if (length(truth) != length(calls)) stop("Length mismatch in calc_sens_fdr: truth=", length(truth), " calls=", length(calls))
  tp <- sum(truth & calls)
  fp <- sum(!truth & calls)
  fn <- sum(truth & !calls)
  list(tp = tp, fp = fp, fn = fn, n_disc = tp + fp, sens = if ((tp + fn) == 0) NA_real_ else tp / (tp + fn), fdr = if ((tp + fp) == 0) 0 else fp / (tp + fp))
}

align_calls_to_taxa <- function(calls, taxa_all, label = "method_call") {
  out <- setNames(rep(FALSE, length(taxa_all)), taxa_all)
  if (is.null(calls)) return(out)
  nm <- names(calls)
  calls <- as.logical(unname(calls))
  names(calls) <- nm
  if (!is.null(nm) && any(nzchar(nm))) {
    idx <- match(nm, taxa_all)
    keep <- !is.na(idx)
    if (any(keep)) out[idx[keep]] <- calls[keep]
    return(out)
  }
  if (length(calls) != length(taxa_all)) stop(label, " returned ", length(calls), " calls but expected ", length(taxa_all), ".")
  setNames(calls, taxa_all)
}

finalise_calls <- function(calls, taxa_cols, label) {
  out <- align_calls_to_taxa(calls, taxa_cols, label)
  out[is.na(out)] <- FALSE
  out
}

pick_first_col <- function(tab, candidates) {
  hit <- intersect(candidates, names(tab))
  if (length(hit)) hit[[1]] else NA_character_
}

extract_calls_from_table <- function(tbl, taxa_names, alpha = 0.05, effect_var = "effect_var", return_details = FALSE) {
  if (is.null(tbl) || nrow(tbl) == 0) {
    out <- setNames(rep(FALSE, length(taxa_names)), taxa_names)
    return(if (return_details) list(calls = out, tested_taxa = character(0)) else out)
  }
  tab <- as_tibble(tbl)
  rn <- rownames(as.data.frame(tbl))
  has_any_taxa_col <- any(c("taxon", "feature", "taxa", "otu", "OTU", "ASV", "id") %in% names(tab))
  if (length(rn) == nrow(tab) && !has_any_taxa_col) tab <- tab %>% mutate(taxon = rn)
  feature_col <- pick_first_col(tab, c("taxon", "feature", "taxa", "otu", "OTU", "ASV", "id"))
  effect_col <- pick_first_col(tab, c("metadata", "coef", "term", "variable", "name"))
  if (!is.na(effect_col)) {
    ef <- as.character(tab[[effect_col]])
    keep <- grepl(paste0("^", effect_var, "$"), ef) | grepl(effect_var, ef, fixed = TRUE)
    if (!any(keep) && effect_var != "cont_cov") keep <- grepl("^cont_cov$", ef) | grepl("cont_cov", ef, fixed = TRUE)
    if (!any(keep) && effect_var != "group") keep <- grepl("^group", ef) | grepl("group$", ef) | grepl("group", ef, fixed = TRUE)
    if (any(keep)) tab <- tab[keep, , drop = FALSE]
  }
  taxa <- if (!is.na(feature_col)) as.character(tab[[feature_col]]) else rownames(as.data.frame(tab))
  call_col <- pick_first_col(tab, c("diff", "reject", "significant", "is_significant", "detected"))
  score <- NULL
  if (!is.na(call_col)) {
    calls_sub <- as.logical(tab[[call_col]])
  } else {
    padj_col <- pick_first_col(tab, c("qval_individual", "qval_joint", "qval", "q_value", "padj", "p_adj", "p.adj", "adj_p", "adj.P.Val", "we.eBH", "wi.eBH", "p.val.adj", "p.adjust"))
    p_col <- pick_first_col(tab, c("pval_individual", "pval_joint", "pval", "p_value", "pvalue", "p", "Pr(>|t|)"))
    if (!is.na(padj_col)) score <- suppressWarnings(as.numeric(tab[[padj_col]]))
    else if (!is.na(p_col)) score <- suppressWarnings(as.numeric(tab[[p_col]]))
    calls_sub <- if (is.null(score)) rep(FALSE, nrow(tab)) else !is.na(score) & score < alpha
  }
  valid_taxa <- taxa %in% taxa_names & !is.na(taxa) & nzchar(taxa)
  tested_taxa <- unique(taxa[valid_taxa & if (!is.na(call_col)) !is.na(calls_sub) else if (!is.null(score)) !is.na(score) else TRUE])
  out <- setNames(rep(FALSE, length(taxa_names)), taxa_names)
  keep <- taxa %in% taxa_names
  if (any(keep)) out[taxa[keep]] <- out[taxa[keep]] | ifelse(is.na(calls_sub[keep]), FALSE, calls_sub[keep])
  if (return_details) list(calls = out, tested_taxa = tested_taxa) else out
}

get_standard_and_robust_calls <- function(res_tab, effect_var = "effect_var") {
  nms <- names(res_tab)
  esc <- gsub("([][{}()+*^$|\\\\?.])", "\\\\\\1", effect_var)
  diff_col <- grep(paste0("^diff_", esc), nms, value = TRUE)[1]
  if (is.na(diff_col) || !diff_col %in% nms) stop("Could not find ANCOM-BC2 standard differential call column (diff_", effect_var, "*).")
  standard <- as.logical(res_tab[[diff_col]])
  standard[is.na(standard)] <- FALSE
  robust_col <- grep(paste0("^diff_robust_", esc), nms, value = TRUE)[1]
  pass_col <- grep(paste0("^passed_ss_", esc), nms, value = TRUE)[1]
  robust <- if (!is.na(robust_col) && robust_col %in% nms) {
    as.logical(res_tab[[robust_col]])
  } else if (!is.na(pass_col) && pass_col %in% nms) {
    standard & as.logical(res_tab[[pass_col]])
  } else {
    NULL
  }
  if (!is.null(robust)) robust[is.na(robust)] <- FALSE
  list(standard = standard, robust = robust)
}

safe_require <- function(pkg) requireNamespace(pkg, quietly = TRUE)

run_ancombc2_calls <- function(y_taxa, md, effect_var, p_adj_method, prv_cut, lib_cut, s0_perc) {
  if (!safe_require("ANCOMBC")) stop("Package 'ANCOMBC' is not installed.")
  fit <- ANCOMBC::ancombc2(
    data = y_taxa, taxa_are_rows = TRUE, meta_data = md, fix_formula = effect_var, group = NULL,
    p_adj_method = p_adj_method, prv_cut = prv_cut, lib_cut = lib_cut, s0_perc = s0_perc,
    struc_zero = FALSE, neg_lb = FALSE, alpha = 0.05, n_cl = 1, verbose = FALSE,
    global = FALSE, pairwise = FALSE, dunnet = FALSE, trend = FALSE
  )
  res_df <- fit$res
  if (is.list(res_df) && "res" %in% names(res_df) && is.data.frame(res_df$res)) res_df <- res_df$res
  res_tab <- res_df %>% as_tibble()
  taxa_all <- rownames(y_taxa)
  taxa_ids <- NULL
  best_match_n <- -1L
  try_candidate <- function(candidate) {
    if (is.null(candidate)) return(NULL)
    candidate <- as.character(candidate)
    if (length(candidate) != nrow(res_tab) || !any(!is.na(candidate) & nzchar(candidate))) return(NULL)
    list(vals = candidate, m = sum(candidate %in% taxa_all, na.rm = TRUE))
  }
  for (cand in list(if (!is.null(fit$feature_table)) rownames(fit$feature_table) else NULL, if ("taxon" %in% names(res_tab)) as.character(res_tab$taxon) else NULL, rownames(res_df))) {
    hit <- try_candidate(cand)
    if (!is.null(hit) && hit$m > best_match_n) {
      taxa_ids <- hit$vals
      best_match_n <- hit$m
    }
  }
  if (is.null(taxa_ids)) stop("ANCOM-BC2: could not determine taxa IDs for result alignment.")
  res_tab$taxon <- taxa_ids
  calls <- get_standard_and_robust_calls(res_tab, effect_var = effect_var)
  list(
    standard = align_calls_to_taxa(setNames(calls$standard, taxa_ids), taxa_all, "ancombc2_standard"),
    robust = if (is.null(calls$robust)) NULL else align_calls_to_taxa(setNames(calls$robust, taxa_ids), taxa_all, "ancombc2_robust"),
    tested_taxa_standard = unique(taxa_ids),
    tested_taxa_robust = unique(taxa_ids)
  )
}

run_maaslin_calls <- function(y_samples, md, taxa_cols, effect_var, sample_loads = NULL) {
  tmp_out <- tempfile(pattern = "maaslin_")
  dir.create(tmp_out, recursive = TRUE, showWarnings = FALSE)
  if (safe_require("maaslin3")) {
    md2 <- md[, c("sample", effect_var), drop = FALSE]
    rownames(md2) <- md2$sample
    md2$sample <- NULL
    args <- list(
      input_data = as.data.frame(y_samples), input_metadata = md2, output = tmp_out,
      formula = paste("~", effect_var), normalization = "TSS", transform = "LOG",
      min_prevalence = 0, correction = "BH", warn_prevalence = FALSE,
      cores = 1, plot_summary_plot = FALSE, plot_associations = FALSE
    )
    if (!is.null(sample_loads)) {
      args$unscaled_abundance <- data.frame(total = as.numeric(sample_loads), row.names = rownames(md2), check.names = FALSE)
      args$median_comparison_abundance <- FALSE
    }
    fit <- do.call(maaslin3::maaslin3, args)
    out_file <- file.path(tmp_out, "all_results.tsv")
    if (!file.exists(out_file)) stop("maaslin3 did not produce all_results.tsv with feature-level results.")
    det <- extract_calls_from_table(readr::read_tsv(out_file, show_col_types = FALSE), taxa_cols, 0.05, effect_var, TRUE)
    return(list(calls = align_calls_to_taxa(det$calls, taxa_cols, "maaslin3_standard"), tested_taxa = det$tested_taxa))
  }
  stop("Package 'maaslin3' is not installed.")
}

run_aldex_calls <- function(y_taxa, md, taxa_cols, effect_var, sample_loads = NULL, gamma = NULL) {
  if (!safe_require("ALDEx3")) stop("Package 'ALDEx3' is not installed.")
  X <- stats::model.matrix(stats::reformulate(effect_var), data = md)
  rownames(X) <- rownames(md)
  X <- X[colnames(y_taxa), , drop = FALSE]
  x_mat <- t(X)
  scale_arg <- ALDEx3::clr.sm
  if (!is.null(sample_loads)) {
    sample_loads <- as.numeric(sample_loads[colnames(y_taxa)])
    if (any(!is.finite(sample_loads)) || any(sample_loads <= 0)) stop("ALDEx3 requires positive finite supplied total microbial load values for every sample.")
    nsample <- 1000L
    scale_arg <- matrix(rep(log2(sample_loads), nsample), nrow = length(sample_loads), ncol = nsample)
  } else {
    nsample <- 1000L
  }
  args <- list(
    Y = y_taxa, X = x_mat, method = "lm", nsample = nsample, n.cores = 1,
    p.adjust.method = "BH", scale = scale_arg, return.pars = c("X", "estimate", "std.error", "p.val", "p.val.adj")
  )
  if (!is.null(gamma)) args$gamma <- gamma
  fit <- do.call(ALDEx3::aldex, args)
  p_adj <- fit$p.val.adj
  if (is.null(p_adj) || !nrow(p_adj)) stop("ALDEx3 returned no adjusted p-value matrix.")
  ridx <- grep(paste0("^", effect_var, "$"), rownames(p_adj))
  if (!length(ridx)) ridx <- grep(effect_var, rownames(p_adj), fixed = TRUE)
  if (!length(ridx)) ridx <- setdiff(seq_len(nrow(p_adj)), 1L)
  if (!length(ridx)) stop("Could not identify ALDEx3 effect row for ", effect_var, ".")
  padj_vec <- setNames(as.numeric(p_adj[ridx[1], ]), colnames(p_adj))
  tab <- tibble(feature = names(padj_vec), p.val.adj = padj_vec, metadata = effect_var)
  det <- extract_calls_from_table(tab, taxa_cols, 0.05, effect_var, TRUE)
  list(calls = align_calls_to_taxa(det$calls, taxa_cols, "aldex3_standard"), tested_taxa = det$tested_taxa)
}

run_locom_calls <- function(y_taxa, md, taxa_cols, effect_var) {
  if (!safe_require("LOCOM")) stop("Package 'LOCOM' is not installed.")
  y_val <- as.numeric(md[[effect_var]])
  # Use permutation test for binary outcomes; asymptotic test for continuous covariates
  is_binary <- length(unique(y_val)) == 2L && all(y_val %in% c(0, 1))
  otu <- t(y_taxa)
  keep <- colMeans(otu > 0) >= 0.2
  otu <- otu[, keep, drop = FALSE]
  fit <- LOCOM::locom(otu.table = otu, Y = y_val, seed = 1, n.cores = 1, n.perm.max = 1000, permute = is_binary, verbose = FALSE)
  qmat <- if (is_binary) fit$q.otu else (fit$q.otu.asymptotic %||% fit$q.otu)
  if (is.null(qmat)) stop("LOCOM returned no q-value matrix.")
  qvec <- as.numeric(qmat[1, ])
  names(qvec) <- colnames(qmat)
  tab <- tibble(taxon = names(qvec), padj = qvec)
  det <- extract_calls_from_table(tab, taxa_cols, 0.05, effect_var, TRUE)
  list(calls = align_calls_to_taxa(det$calls, taxa_cols, "locom_standard"), tested_taxa = det$tested_taxa)
}

run_linda_calls <- function(y_taxa, md, taxa_cols, effect_var) {
  if (!safe_require("MicrobiomeStat")) stop("Package 'MicrobiomeStat' is not installed.")
  fit <- MicrobiomeStat::linda(feature.dat = y_taxa, meta.dat = md, formula = paste("~", effect_var), feature.dat.type = "count", prev.filter = 0, p.adj.method = "BH", alpha = 0.05, verbose = FALSE)
  tab <- NULL
  if (is.list(fit) && !is.null(fit$output) && is.list(fit$output)) {
    tab <- fit$output[[1]]
    if (!is.null(tab)) tab <- as_tibble(tab, rownames = "taxon")
  }
  if (is.list(fit) && !is.null(fit$res)) tab <- fit$res
  det <- extract_calls_from_table(tab, taxa_cols, 0.05, effect_var, TRUE)
  list(calls = align_calls_to_taxa(det$calls, taxa_cols, "linda_standard"), tested_taxa = det$tested_taxa)
}

run_wilcoxon_calls <- function(y_samples, taxa_cols, effect_values, sample_loads = NULL, p_adj_method = "BH") {
  effect_values <- as.numeric(effect_values)
  if (anyNA(effect_values)) stop("Wilcoxon testing requires non-missing effect values.")
  if (!all(effect_values %in% c(0, 1))) stop("Wilcoxon testing is only defined here for binary comparisons; continuous covariates are not supported.")
  if (length(unique(effect_values)) != 2) stop("Wilcoxon testing requires both comparison groups to be present.")
  if (!is.null(sample_loads)) {
    sample_loads <- as.numeric(sample_loads)
    if (length(sample_loads) != nrow(y_samples)) stop("Sample load vector length does not match number of samples.")
    if (anyNA(sample_loads) || any(!is.finite(sample_loads)) || any(sample_loads <= 0)) stop("Wilcoxon sample-load scaling requires positive, finite sample loads.")
    y_samples <- sweep(y_samples, 1, sample_loads, "*")
  }
  g0 <- which(effect_values == 0)
  g1 <- which(effect_values == 1)
  pvals <- vapply(seq_along(taxa_cols), function(j) {
    tryCatch(stats::wilcox.test(y_samples[g1, j], y_samples[g0, j], exact = FALSE)$p.value, error = function(e) NA_real_)
  }, numeric(1))
  padj <- stats::p.adjust(pvals, method = p_adj_method)
  list(calls = setNames(!is.na(padj) & padj < 0.05, taxa_cols), tested_taxa = taxa_cols[!is.na(padj)])
}

job_stems <- function(job) {
  list(
    results = job$results_stem %||% "multimethod_results",
    failures = job$failures_stem %||% "multimethod_failures"
  )
}

build_result_row <- function(stem, fname, mrow, cmp, method_name, d, truth, n_samples, metrics, n_tested, n_unmatched) {
  tibble(
    dataset_id = stem, file = fname, scenario_name = as.character(mrow$scenario_name[[1]]),
    comparison_name = cmp$comparison_name, reference_group = cmp$reference_group, target_group = cmp$target_group,
    method = method_name, spikeFraction = as.numeric(mrow$spike_fraction[[1]]),
    sampleSizePerGroup = as.numeric(mrow$sample_size_per_group[[1]]), rep = as.numeric(mrow$replicate[[1]]),
    nGroups = as.numeric(mrow$n_groups[[1]]), nSamplesTotal = as.numeric(mrow$n_samples_total[[1]]),
    nSamplesCompared = n_samples, nTaxa = d, nTrue = sum(truth), nTaxaTested = n_tested,
    nTaxaFiltered = d - n_tested, nTaxaUnmatched = n_unmatched, nDiscoveries = metrics$n_disc,
    TP = metrics$tp, FP = metrics$fp, FN = metrics$fn, Sensitivity = metrics$sens, FDR = metrics$fdr
  )
}

run_method <- function(job, method, y_taxa, y_samples, md, taxa_cols, cmp) {
  sample_loads <- if ("total_microbial_load" %in% names(md)) setNames(md$total_microbial_load, md$sample) else (cmp$sample_loads %||% NULL)
  if (method == "ancombc2") return(run_ancombc2_calls(y_taxa, md, "effect_var", job$p_adj_method, job$prv_cut, job$lib_cut, job$s0_perc))
  if (method == "maaslin3") return(with(run_maaslin_calls(y_samples, md, taxa_cols, "effect_var", sample_loads), list(standard = calls, robust = NULL, tested_taxa_standard = tested_taxa, tested_taxa_robust = NULL)))
  if (method == "aldex3") return(with(run_aldex_calls(y_taxa, md, taxa_cols, "effect_var", sample_loads, job$aldex_gamma %||% NULL), list(standard = calls, robust = NULL, tested_taxa_standard = tested_taxa, tested_taxa_robust = NULL)))
  if (method == "locom") return(with(run_locom_calls(y_taxa, md, taxa_cols, "effect_var"), list(standard = calls, robust = NULL, tested_taxa_standard = tested_taxa, tested_taxa_robust = NULL)))
  if (method == "linda") return(with(run_linda_calls(y_taxa, md, taxa_cols, "effect_var"), list(standard = calls, robust = NULL, tested_taxa_standard = tested_taxa, tested_taxa_robust = NULL)))
  if (method == "wilcoxon") {
    # Wilcoxon is only defined for binary comparisons; silently skip continuous covariate datasets
    if (!all(as.numeric(cmp$effect_values) %in% c(0, 1))) return(NULL)
    res <- run_wilcoxon_calls(y_samples, taxa_cols, cmp$effect_values, sample_loads, "BH")
    return(list(standard = res$calls, robust = NULL, tested_taxa_standard = res$tested_taxa, tested_taxa_robust = NULL))
  }
  stop("Unknown method: ", method)
}

run_job <- function(project_dir, job) {
  out_dir <- Sys.getenv("OUT_DIR", unset = "")
  if (!nzchar(out_dir)) out_dir <- file.path(project_dir, job$out_subdir)
  dir.create(out_dir, recursive = TRUE, showWarnings = FALSE)
  stems <- job_stems(job)
  fail_csv_path <- file.path(out_dir, paste0(stems$failures, ".csv"))
  fail_progress_csv_path <- file.path(out_dir, paste0(stems$failures, "_progress.csv"))
  data_dir <- file.path(project_dir, "simulated_count_tables")
  meta_file <- file.path(data_dir, "biased_simulations_metadata.csv")
  if (!dir.exists(data_dir)) stop("Missing directory: ", data_dir)
  if (!file.exists(meta_file)) stop("Missing metadata CSV: ", meta_file)
  meta <- readr::read_csv(meta_file, show_col_types = FALSE)
  req_cols <- c("dataset_id", "scenario_name", "output_csv", "spike_fraction", "sample_size_per_group", "replicate", "n_groups", "n_samples_total")
  if (!all(req_cols %in% names(meta))) stop("Metadata is missing required columns: ", paste(setdiff(req_cols, names(meta)), collapse = ", "))
  meta <- meta %>% mutate(dataset_path = vapply(output_csv, resolve_dataset_path, character(1), meta_file = meta_file))
  missing_paths <- meta$dataset_path[!file.exists(meta$dataset_path)]
  if (length(missing_paths)) stop("Metadata references missing count tables, e.g. ", missing_paths[[1]])
  max_files <- suppressWarnings(as.numeric(Sys.getenv("MAX_FILES", unset = "Inf")))
  if (is.finite(max_files)) meta <- meta[seq_len(min(nrow(meta), max_files)), , drop = FALSE]
  checkpoint_every <- suppressWarnings(as.integer(Sys.getenv("CHECKPOINT_EVERY", unset = "100")))
  if (is.na(checkpoint_every)) checkpoint_every <- 5L
  latest_checkpoint <- find_latest_checkpoint(out_dir, stems$results)
  checkpoint_done <- 0L
  existing_results <- existing_failures <- NULL
  if (!is.null(latest_checkpoint) && file.exists(latest_checkpoint)) {
    checkpoint_done <- min(nrow(meta), suppressWarnings(as.integer(sub(".*_checkpoint_([0-9]{4})\\.csv$", "\\1", basename(latest_checkpoint)))))
    existing_results <- readr::read_csv(latest_checkpoint, show_col_types = FALSE)
    if (file.exists(fail_progress_csv_path)) existing_failures <- readr::read_csv(fail_progress_csv_path, show_col_types = FALSE)
  }
  methods <- job$methods
  methods_env <- Sys.getenv("METHODS", unset = "")
  if (nzchar(methods_env)) methods <- intersect(methods, trimws(strsplit(methods_env, ",", fixed = TRUE)[[1]]))
  res_rows <- vector("list", max(1, nrow(meta) * max(1, length(methods)) * 4))
  fail_rows <- vector("list", max(1, nrow(meta) * max(1, length(methods)) * 4))
  row_idx <- fail_idx <- 0L
  if (!is.null(existing_results) && nrow(existing_results)) { row_idx <- nrow(existing_results); res_rows[seq_len(row_idx)] <- split(existing_results, seq_len(nrow(existing_results))) }
  if (!is.null(existing_failures) && nrow(existing_failures)) { fail_idx <- nrow(existing_failures); fail_rows[seq_len(fail_idx)] <- split(existing_failures, seq_len(nrow(existing_failures))) }
  n_done <- checkpoint_done
  if (n_done < nrow(meta)) for (i in (n_done + 1L):nrow(meta)) {
    mrow <- meta[i, , drop = FALSE]
    fpath <- mrow$dataset_path[[1]]
    fname <- basename(fpath)
    stem <- sub("\\.csv$", "", fname)
    message(sprintf("[%s %d/%d] %s", job$id, i, nrow(meta), fname))
    dat <- readr::read_csv(fpath, show_col_types = FALSE)
    taxa_cols <- get_taxa_columns(dat)
    if (!length(taxa_cols)) stop("No numeric taxa columns found in file: ", fname)
    d <- length(taxa_cols)
    comparisons <- build_dataset_comparisons(mrow, dat, taxa_cols, isTRUE(job$sample_load_mode))
    for (cmp in comparisons) {
      dat_cmp <- dat[cmp$row_idx, , drop = FALSE]
      sample_ids <- paste0(stem, "_S", seq_len(nrow(dat_cmp)))
      y_samples <- as.matrix(dat_cmp[, taxa_cols, drop = FALSE])
      storage.mode(y_samples) <- "numeric"
      colnames(y_samples) <- taxa_cols
      rownames(y_samples) <- sample_ids
      y_taxa <- t(y_samples)
      colnames(y_taxa) <- rownames(y_samples)
      rownames(y_taxa) <- taxa_cols
      md <- tibble(sample = sample_ids, effect_var = as.numeric(cmp$effect_values)) %>% as.data.frame()
      if (isTRUE(job$sample_load_mode)) md$total_microbial_load <- as.numeric(cmp$sample_loads)
      rownames(md) <- md$sample
      truth <- setNames(as.logical(cmp$truth), taxa_cols)
      for (method in methods) {
        message("  - ", cmp$comparison_name, " / ", method)
        result <- tryCatch(run_method(job, method, y_taxa, y_samples, md, taxa_cols, cmp), error = function(e) {
          fail_idx <<- fail_idx + 1L
          fail_rows[[fail_idx]] <<- tibble(dataset_id = stem, file = fname, scenario_name = as.character(mrow$scenario_name[[1]]), comparison_name = cmp$comparison_name, method = method, error = conditionMessage(e))
          NULL
        })
        if (is.null(result)) next
        calls_std <- finalise_calls(result$standard, taxa_cols, paste0(method, "_standard_final"))
        tested_std_raw <- unique(as.character(result$tested_taxa_standard))
        tested_std_raw <- tested_std_raw[!is.na(tested_std_raw) & nzchar(tested_std_raw)]
        tested_std <- unique(intersect(tested_std_raw, taxa_cols))
        if (length(tested_std_raw) && !length(tested_std)) stop("Taxa mapping failed for ", method, " in ", fname, " / ", cmp$comparison_name)
        metrics_std <- calc_sens_fdr(truth, calls_std)
        row_idx <- row_idx + 1L
        res_rows[[row_idx]] <- build_result_row(stem, fname, mrow, cmp, switch(method, ancombc2 = "ancombc2_standard", wilcoxon = "wilcoxon_tss", method), d, truth, nrow(y_samples), metrics_std, length(tested_std), length(setdiff(tested_std_raw, taxa_cols)))
        if (method == "ancombc2") {
          if (is.null(result$robust)) stop("ANCOM-BC2 did not return SS-filtered (robust) calls.")
          calls_rob <- finalise_calls(result$robust, taxa_cols, "ancombc2_robust_final")
          tested_rob_raw <- unique(as.character(result$tested_taxa_robust))
          tested_rob_raw <- tested_rob_raw[!is.na(tested_rob_raw) & nzchar(tested_rob_raw)]
          tested_rob <- unique(intersect(tested_rob_raw, taxa_cols))
          metrics_rob <- calc_sens_fdr(truth, calls_rob)
          row_idx <- row_idx + 1L
          res_rows[[row_idx]] <- build_result_row(stem, fname, mrow, cmp, "ancombc2_ss_filtered", d, truth, nrow(y_samples), metrics_rob, length(tested_rob), length(setdiff(tested_rob_raw, taxa_cols)))
        }
      }
    }
    n_done <- n_done + 1L
    if (checkpoint_every > 0 && n_done %% checkpoint_every == 0) {
      partial_results <- bind_rows(res_rows) %>% arrange(scenario_name, comparison_name, method, spikeFraction, sampleSizePerGroup, rep)
      readr::write_csv(partial_results, file.path(out_dir, sprintf("%s_checkpoint_%04d.csv", stems$results, n_done)))
      readr::write_csv(partial_results, file.path(out_dir, paste0(stems$results, "_progress.csv")))
      if (fail_idx > 0) readr::write_csv(bind_rows(fail_rows), fail_progress_csv_path)
    }
  }
  results <- bind_rows(res_rows)
  if (!nrow(results)) stop("No valid results were produced.")
  results <- results %>% arrange(scenario_name, comparison_name, method, spikeFraction, sampleSizePerGroup, rep)
  readr::write_csv(results, file.path(out_dir, paste0(stems$results, ".csv")))
  if (fail_idx > 0) readr::write_csv(bind_rows(fail_rows), fail_csv_path)
}

run_project_jobs <- function(project_dir, jobs) {
  filter <- Sys.getenv("JOB_FILTER", unset = "")
  if (nzchar(filter)) jobs <- jobs[vapply(jobs, function(x) x$id %in% trimws(strsplit(filter, ",", fixed = TRUE)[[1]]), logical(1))]
  if (!length(jobs)) stop("No jobs selected.")
  for (job in jobs) {
    message("Running job: ", job$id, " -> ", job$out_subdir)
    job$p_adj_method <- job$p_adj_method %||% "holm"
    job$prv_cut <- job$prv_cut %||% 0.10
    job$lib_cut <- job$lib_cut %||% 0
    job$s0_perc <- job$s0_perc %||% 0.05
    run_job(project_dir, job)
  }
}
```
