## Supplementary Code for "Bias-mitigated microbiome inference refines coronary artery disease signature": aggregate_hpc_dataset_results.m

```
function Results = aggregate_hpc_dataset_results(projectRoot)
    if nargin < 1 || isempty(projectRoot)
        projectRoot = pwd;
    end

    resultDir = fullfile(projectRoot, "simulated_count_tables_analysis_hpc", "dataset_results");
    outCsv = fullfile(projectRoot, "simulated_count_tables_analysis_hpc", "biased_simulations_fdr_sensitivity_results.csv");
    files = dir(fullfile(resultDir, "*.csv"));
    if isempty(files)
        error("No per-dataset result CSVs found in %s", resultDir);
    end

    parts = cell(numel(files), 1);
    for i = 1:numel(files)
        parts{i} = readtable(fullfile(files(i).folder, files(i).name), 'TextType', 'string');
    end
    Results = vertcat(parts{:});
    Results = sortrows(Results, {'scenario_name','comparison_name','spikeFraction','sampleSizePerGroup','rep'});
    writetable(Results, outCsv);
    fprintf('Aggregated %d per-dataset result files into %s\n', numel(files), outCsv);
end
```
