## Supplementary Code for "Bias-mitigated microbiome inference refines coronary artery disease signature": als_fit_orthog_hpc.m

```
function out = als_fit_orthog_hpc(Y, X, metadata, prescribed_size, prescribed_contam)
    maxIter = 1000;
    tol = 10;
    epsS = 1e-12;

    [n, d] = size(Y);
    good = any(Y ~= 0, 2);

    if ~any(good)
        out.B = zeros(size(X,2), d);
        out.S = zeros(n,1);
        out.Ghat = out.B(2:end,:);
        out.loss = 0;
        out.iters = 0;
        return
    end

    Yr = Y(good,:);
    Xr = X(good,:);

    sampleBiasAll = parse_metadata_vector_hpc(metadata, "sample_bias_factors");
    contamAll = parse_metadata_vector_hpc(metadata, "contam_taxa_bias_factors");

    if prescribed_size == 1 && prescribed_contam == 1
        Sr = sampleBiasAll(good);
        Aj = contamAll(:)';
        Aj = Aj(1:d);
        B = zeros(size(Xr,2), d);
        B(1,:) = Aj;
        if size(Xr,2) > 1
            B(2:end,:) = Xr(:,2:end) \ ((Yr ./ Sr) - Aj);
        end
    elseif prescribed_size == 1
        Sr = sampleBiasAll(good);
        B = Xr \ (Yr ./ Sr);
    elseif prescribed_contam == 1
        Aj = contamAll(:)';
        Aj = Aj(1:d);
        B = zeros(size(Xr,2), d);
        B(1,:) = Aj;
        Sr = max(mean(Yr,2), epsS);
        Sr = Sr ./ mean(Sr);
        if size(Xr,2) > 1
            B(2:end,:) = Xr(:,2:end) \ ((Yr ./ Sr) - Aj);
        end
    else
        Sr = max(mean(Yr,2), epsS);
        Sr = Sr ./ mean(Sr);
        B = Xr \ (Yr ./ Sr);
    end

    lossPrev = inf;
    for it = 1:maxIter
        Yt = Yr ./ Sr;
        B = Xr \ Yt;
        Mu = Xr * B;
        num = sum(Mu .* Yr, 2);
        den = sum(Mu .* Mu, 2);
        Sr = num ./ max(den, epsS);
        Sr = max(Sr, epsS);
        Sr = Sr ./ mean(Sr);
        lr = log(max(Sr, epsS));
        beta_s = Xr \ lr;
        lr = lr - Xr(:,2:end) * beta_s(2:end);
        Sr = exp(lr);
        Sr = Sr ./ mean(Sr);
        R = Yr - Sr .* Mu;
        loss = sum(R(:).^2);
        if abs(lossPrev - loss) / max(lossPrev, 1) < tol
            break
        end
        lossPrev = loss;
    end

    S = zeros(n,1);
    S(good) = Sr;

    out.B = B;
    out.S = S;
    out.Ghat = B(2:end,:);
    out.loss = loss;
    out.iters = it;
    out.good = good;
end
```
