## Supplementary Code for "Bias-mitigated microbiome inference refines coronary artery disease signature": ALS_pair_boot_biased_metadata_orthog_hpc.m

```
function ALS_pair_boot_biased_metadata_orthog_hpc(projectRoot, alpha, maxBoot, B_override, useParallel, numWorkers, prescribed_size, prescribed_contam)
% HPC worker entrypoint. Claims datasets via lock files, processes them one
% at a time, and writes one result CSV per dataset so multiple jobs can run
% concurrently without clobbering a shared results file.

    if nargin < 1 || isempty(projectRoot)
        projectRoot = pwd;
    end
    if nargin < 2 || isempty(alpha), alpha = 0.05; end
    if nargin < 3 || isempty(maxBoot), maxBoot = inf; end
    if nargin < 4, B_override = []; end
    if nargin < 5 || isempty(useParallel), useParallel = true; end
    if nargin < 6 || isempty(numWorkers), numWorkers = 16; end
    if nargin < 7 || isempty(prescribed_size), prescribed_size = 0; end
    if nargin < 8 || isempty(prescribed_contam), prescribed_contam = 0; end

    dataDir = fullfile(projectRoot, "simulated_count_tables");
    metaFile = fullfile(dataDir, "biased_simulations_metadata.csv");
    outDir = fullfile(projectRoot, "simulated_count_tables_analysis_hpc");
    datasetResultDir = fullfile(outDir, "dataset_results");
    lockDir = fullfile(outDir, "locks");
    tmpDir = projectRoot;

    ensure_folder_hpc(outDir);
    ensure_folder_hpc(datasetResultDir);
    ensure_folder_hpc(lockDir);

    if useParallel && numWorkers > 1
        p = gcp('nocreate');
        if isempty(p) || p.NumWorkers ~= numWorkers
            if ~isempty(p), delete(p); end
            parpool('local', numWorkers);
        end
    end

    Tmeta = read_metadata_table_hpc(metaFile);
    requiredCols = ["dataset_id", "output_csv", "scenario_name"];
    if ~all(ismember(requiredCols, string(Tmeta.Properties.VariableNames)))
        error("Metadata must include dataset_id, output_csv, and scenario_name columns.");
    end

    fprintf('HPC worker starting in %s\n', projectRoot);
    while true
        [metaRow, rowIdx, lockPath, resultPath] = claim_next_dataset_lock_hpc(Tmeta, datasetResultDir, lockDir);
        if isempty(rowIdx)
            fprintf('No unlocked datasets remain. Worker exiting.\n');
            break
        end

        cleanupLock = onCleanup(@() release_lock_hpc(lockPath));
        datasetId = string(metaRow.dataset_id);
        fprintf('Claimed dataset %d/%d: %s\n', rowIdx, height(Tmeta), datasetId);

        try
            datasetResults = process_dataset_hpc(metaRow, rowIdx, alpha, maxBoot, B_override, useParallel, prescribed_size, prescribed_contam, tmpDir);
            writetable(datasetResults, resultPath);
            fprintf('Completed %s -> %s\n', datasetId, resultPath);
        catch ME
            errPath = fullfile(lockDir, datasetId + ".error.txt");
            fid = fopen(errPath, 'w');
            if fid ~= -1
                fprintf(fid, '%s\n', getReport(ME, 'extended'));
                fclose(fid);
            end
            rethrow(ME)
        end

        clear cleanupLock
    end
end
```
