## Supplementary Code for "Bias-mitigated microbiome inference refines coronary artery disease signature": build_comparisons_hpc.m

```
function comparisons = build_comparisons_hpc(metaRow, Y, groupVec, contVec, d)
    scenario = string(metaRow.scenario_name);
    betaByGroup = zeros(3, d);
    for g = 1:3
        [idx, beta] = parse_index_and_beta_hpc(metaRow, g);
        valid = idx >= 1 & idx <= d;
        betaByGroup(g, idx(valid)) = beta(valid);
    end

    switch scenario
        case "binary_covariate"
            mask = ismember(groupVec, [1 2]);
            comparisons = struct( ...
                'name', "group2_vs_group1", ...
                'referenceGroup', 1, ...
                'targetGroup', 2, ...
                'rowIdx', find(mask), ...
                'Y', Y(mask,:), ...
                'X', [ones(sum(mask),1), double(groupVec(mask) == 2)], ...
                'Gtrue', abs(betaByGroup(2,:) - betaByGroup(1,:)) > 0);

        case "continuous_covariate"
            if isempty(contVec)
                comparisons = struct([]);
                return
            end
            active = any(abs(betaByGroup) > 0, 1);
            comparisons = struct( ...
                'name', "continuous_effect", ...
                'referenceGroup', NaN, ...
                'targetGroup', NaN, ...
                'rowIdx', (1:size(Y,1))', ...
                'Y', Y, ...
                'X', [ones(size(Y,1),1), contVec(:)], ...
                'Gtrue', active);

        case "three_group_spikein"
            pairList = [1 2; 1 3; 2 3];
            comparisons = repmat(struct( ...
                'name', "", ...
                'referenceGroup', NaN, ...
                'targetGroup', NaN, ...
                'rowIdx', [], ...
                'Y', [], ...
                'X', [], ...
                'Gtrue', []), 1, size(pairList,1));

            for i = 1:size(pairList,1)
                ref = pairList(i,1);
                tgt = pairList(i,2);
                mask = ismember(groupVec, [ref tgt]);
                comparisons(i).name = sprintf('group%d_vs_group%d', tgt, ref);
                comparisons(i).referenceGroup = ref;
                comparisons(i).targetGroup = tgt;
                comparisons(i).rowIdx = find(mask);
                comparisons(i).Y = Y(mask,:);
                comparisons(i).X = [ones(sum(mask),1), double(groupVec(mask) == tgt)];
                comparisons(i).Gtrue = abs(betaByGroup(tgt,:) - betaByGroup(ref,:)) > 0;
            end

        otherwise
            comparisons = struct([]);
    end
end
```
