## Supplementary Code for "Bias-mitigated microbiome inference refines coronary artery disease signature": claim_next_dataset_lock_hpc.m

```
function [metaRow, rowIdx, lockPath, resultPath] = claim_next_dataset_lock_hpc(Tmeta, datasetResultDir, lockDir)
    metaRow = table();
    rowIdx = [];
    lockPath = "";
    resultPath = "";

    for i = 1:height(Tmeta)
        datasetId = string(Tmeta.dataset_id(i));
        resultPathTry = fullfile(datasetResultDir, datasetId + ".csv");
        if isfile(resultPathTry)
            continue
        end

        lockPathTry = fullfile(lockDir, datasetId + ".lock");
        if isfile(lockPathTry)
            continue
        end
        fid = fopen(lockPathTry, 'w');
        if fid == -1
            continue
        end
        fprintf(fid, 'pid=%d\n', feature('getpid'));
        fclose(fid);

        metaRow = Tmeta(i,:);
        rowIdx = i;
        lockPath = lockPathTry;
        resultPath = resultPathTry;
        return
    end
end
```
