## Supplementary Code for "Bias-mitigated microbiome inference refines coronary artery disease signature": draw_bootstrap_idx_hpc.m

```
function idx = draw_bootstrap_idx_hpc(N, rowsByGroup, isBinary, seed, b)
    rng(seed + b, 'twister');
    if isBinary
        idx = zeros(N, 1);
        pos = 1;
        for k = 1:numel(rowsByGroup)
            rows = rowsByGroup{k};
            nk = numel(rows);
            draw = rows(randi(nk, nk, 1));
            idx(pos:pos+nk-1) = draw;
            pos = pos + nk;
        end
    else
        idx = randi(N, N, 1);
    end
end
```
