## Supplementary Code for "Bias-mitigated microbiome inference refines coronary artery disease signature": extract_design_inputs_hpc.m

```
function [Y, groupVec, contVec] = extract_design_inputs_hpc(T, datasetId, scenarioName)
    if nargin < 2 || strlength(string(datasetId)) == 0
        datasetId = "unknown_dataset";
    end
    if nargin < 3 || strlength(string(scenarioName)) == 0
        scenarioName = "";
    end

    varNames = string(T.Properties.VariableNames);
    metaCols = ["synthetic_sample_id", "source_sample_id", "group", "continuous_covariate"];
    isNumericVar = varfun(@isnumeric, T, 'OutputFormat', 'uniform');
    taxaMask = isNumericVar & ~ismember(varNames, metaCols) & ~startsWith(varNames, "ExtraVar");
    Y = double(table2array(T(:, taxaMask)));

    if ismember("group", varNames)
        groupRaw = strtrim(string(T.group));
        groupClean = regexprep(groupRaw, '[^0-9\.\-\+]', '');
        groupVec = double(str2double(groupClean));
    else
        groupVec = ones(height(T), 1);
    end

    if ismember("continuous_covariate", varNames)
        contRaw = strtrim(string(T.continuous_covariate));
        contClean = regexprep(contRaw, '[^0-9eE\.\-\+]', '');
        contVec = double(str2double(contClean));
    else
        contVec = [];
    end

    switch string(scenarioName)
        case {"binary_covariate", "three_group_spikein"}
            if ~ismember("group", varNames)
                error("Dataset %s is missing required 'group' column. Imported columns: %s", ...
                    datasetId, strjoin(cellstr(varNames), ', '));
            end
            if all(isnan(groupVec))
                error("Dataset %s has a 'group' column but none of its values parsed numerically.", datasetId);
            end

            groupsPresent = unique(groupVec(~isnan(groupVec)))';
            if scenarioName == "binary_covariate"
                if ~all(ismember([1 2], groupsPresent))
                    error("Dataset %s expected groups 1 and 2, but parsed groups were: %s", ...
                        datasetId, join(string(groupsPresent), ','));
                end
            else
                if ~all(ismember([1 2 3], groupsPresent))
                    error("Dataset %s expected groups 1, 2 and 3, but parsed groups were: %s", ...
                        datasetId, join(string(groupsPresent), ','));
                end
            end

        case "continuous_covariate"
            if ~ismember("continuous_covariate", varNames)
                error("Dataset %s is missing required 'continuous_covariate' column. Imported columns: %s", ...
                    datasetId, strjoin(cellstr(varNames), ', '));
            end
            if all(isnan(contVec))
                error("Dataset %s has a 'continuous_covariate' column but none of its values parsed numerically.", datasetId);
            end
    end
end
```
