## Supplementary Code for "Bias-mitigated microbiome inference refines coronary artery disease signature": generate_count_tables.m

```
%% simulate count tables across multiple covariate designs
%% USER SETTINGS
clear
clc
close all
rng(1)

inputCsv = "qmp_full.csv";
flowLoadsCsv = "flow_loads.csv";
outputDir = "simulated_count_tables";
metadataCsv = "biased_simulations_metadata.csv";

sampleSizesPerGroup = [30 50 100 200 300];
nReplicates = 100;
spikeFractions = [0.1 0.5 0.9];
spikeFoldSetSigned = [-10 -5 -2 2 5 10];

taxaKeepFraction = 0.118;
kPosRequired = 8;
allowOverlapSpikes = true;
contaminationFraction = 0.5;

scenarios = [ ...
    struct('name',"binary_covariate",'label',"Binary covariate",'nGroups',2,'usesContinuous',false,'hasSpike',true,'groupSpikeTargets',2)
];

if ~isfolder(outputDir)
    mkdir(outputDir);
end

T = readtable(inputCsv, 'PreserveVariableNames', true, 'TextType', 'string');
varNames = string(T.Properties.VariableNames);

hasSampleId = strcmpi(varNames(1), "sample_id");
if hasSampleId
    sampleIds = string(T{:,1});
    countTable = T(:,2:end);
    taxaNames = varNames(2:end);
else
    sampleIds = "source_" + compose('%05d', (1:height(T))');
    countTable = T;
    taxaNames = varNames;
end

S_i = readtable(flowLoadsCsv, 'PreserveVariableNames', true);
sampleBiasPool = table2array(S_i(:,3));
sampleBiasPool = double(sampleBiasPool(:));
sampleBiasPool = sampleBiasPool ./ mean(sampleBiasPool);

colsums = table2array(sum(countTable, 1));
countTable(:, colsums == 0) = [];
counts = double(table2array(countTable));

keepIdx = selectTaxaToKeep(counts, taxaKeepFraction);
counts = counts(:, keepIdx);
taxaNames = taxaNames(keepIdx);
[nSamples, nTaxa] = size(counts);

nRows = estimateMetadataRows(scenarios, spikeFractions, sampleSizesPerGroup, nReplicates);
metadataVarNames = [ ...
    "dataset_id", "scenario_name", "scenario_label", "output_csv", ...
    "replicate", "spike_fraction", "sample_size_per_group", ...
    "n_groups", "n_samples_total", "n_taxa", ...
    "group_counts", "group_labels", "continuous_covariate", ...
    "spike_taxa_indices_group1", "spike_taxa_indices_group2", "spike_taxa_indices_group3", ...
    "spike_fold_changes_signed_group1", "spike_fold_changes_signed_group2", "spike_fold_changes_signed_group3", ...
    "spike_beta_group1", "spike_beta_group2", "spike_beta_group3", ...
    "taxa_bias_factors", "contam_taxa_bias_factors", "sample_bias_factors", ...
    "source_sample_ids", "notes" ...
];
metadata = table('Size', [nRows, numel(metadataVarNames)], ...
    'VariableTypes', repmat("string", 1, numel(metadataVarNames)), ...
    'VariableNames', metadataVarNames);

row = 0;

for s = 1:numel(scenarios)
    scenario = scenarios(s);

    scenarioSpikeFractions = spikeFractions;

    for i = 1:numel(scenarioSpikeFractions)
        spikeFrac = scenarioSpikeFractions(i);
        nSpiked = max(1, round(spikeFrac * nTaxa));

        for j = 1:numel(sampleSizesPerGroup)
            nPerGroup = sampleSizesPerGroup(j);
            nCurrSamples = scenario.nGroups * nPerGroup;

            for r = 1:nReplicates
                row = row + 1;
                datasetId = sprintf('%s_sf%02d_npg%03d_rep%03d', ...
                    scenario.name, round(100 * spikeFrac), nPerGroup, r);

                sourceIdx = randi(nSamples, nCurrSamples, 1);
                countsSub = counts(sourceIdx, :);
                groupLabelsCurr = makeGroupLabels(scenario.nGroups, nPerGroup);

                sampleFactors = sampleBiasPool(randi(numel(sampleBiasPool), nCurrSamples, 1));
                taxaFactors = unifrnd(0.1, 1.0, 1, nTaxa);
                baselineContam = buildBaselineContamination(counts, nTaxa, contaminationFraction);

                alphaEff = countsSub;
                x = strings(nCurrSamples, 1);

                if scenario.usesContinuous
                    xNumeric = randn(nCurrSamples, 1);
                    x = compose('%.6g', xNumeric);
                end

                spikeInfo = initialiseSpikeInfo();

                if scenario.hasSpike
                    eligibleCols = findEligibleTaxa(countsSub, groupLabelsCurr, kPosRequired);
                    spikeInfo = assignScenarioSpikes( ...
                        scenario, eligibleCols, nSpiked, spikeFoldSetSigned, ...
                        allowOverlapSpikes, x, alphaEff, groupLabelsCurr);
                    alphaEff = spikeInfo.alphaEff;
                end

                inner = repmat(taxaFactors, nCurrSamples, 1) .* alphaEff + repmat(baselineContam, nCurrSamples, 1);
                inner(inner < 0) = 0;
                mu = sampleFactors .* inner;
                simCounts = round(mu);

                outTable = array2table(simCounts, 'VariableNames', cellstr(taxaNames));
                syntheticIds = datasetId + "_sample_" + compose('%05d', (1:nCurrSamples)');
                outTable = addvars(outTable, syntheticIds, string(groupLabelsCurr), ...
                    'Before', 1, 'NewVariableNames', {'synthetic_sample_id', 'group'});
                outTable = addvars(outTable, sampleIds(sourceIdx), 'Before', 3, ...
                    'NewVariableNames', {'source_sample_id'});

                if scenario.usesContinuous
                    outTable = addvars(outTable, x, 'After', 'group', ...
                        'NewVariableNames', {'continuous_covariate'});
                end

                outPath = fullfile(outputDir, datasetId + ".csv");
                writetable(outTable, outPath);

                metadata.dataset_id(row) = datasetId;
                metadata.scenario_name(row) = scenario.name;
                metadata.scenario_label(row) = scenario.label;
                metadata.output_csv(row) = string(outPath);
                metadata.replicate(row) = string(r);
                metadata.spike_fraction(row) = compose('%.6g', spikeFrac);
                metadata.sample_size_per_group(row) = string(nPerGroup);
                metadata.n_groups(row) = string(scenario.nGroups);
                metadata.n_samples_total(row) = string(nCurrSamples);
                metadata.n_taxa(row) = string(nTaxa);
                metadata.group_counts(row) = summariseGroupCounts(groupLabelsCurr);
                metadata.group_labels(row) = serialiseStrings(string(groupLabelsCurr'));
                metadata.continuous_covariate(row) = serialiseStrings(x');
                metadata.spike_taxa_indices_group1(row) = serialiseNumeric(spikeInfo.indices{1});
                metadata.spike_taxa_indices_group2(row) = serialiseNumeric(spikeInfo.indices{2});
                metadata.spike_taxa_indices_group3(row) = serialiseNumeric(spikeInfo.indices{3});
                metadata.spike_fold_changes_signed_group1(row) = serialiseNumeric(spikeInfo.foldChanges{1});
                metadata.spike_fold_changes_signed_group2(row) = serialiseNumeric(spikeInfo.foldChanges{2});
                metadata.spike_fold_changes_signed_group3(row) = serialiseNumeric(spikeInfo.foldChanges{3});
                metadata.spike_beta_group1(row) = serialiseNumeric(spikeInfo.betas{1});
                metadata.spike_beta_group2(row) = serialiseNumeric(spikeInfo.betas{2});
                metadata.spike_beta_group3(row) = serialiseNumeric(spikeInfo.betas{3});
                metadata.taxa_bias_factors(row) = serialiseNumeric(taxaFactors);
                metadata.contam_taxa_bias_factors(row) = serialiseNumeric(baselineContam);
                metadata.sample_bias_factors(row) = serialiseNumeric(sampleFactors');
                metadata.source_sample_ids(row) = serialiseStrings(sampleIds(sourceIdx)');
                metadata.notes(row) = scenarioNotes(scenario);
            end

            fprintf('Completed %s, spike fraction %.2f, sample size per group %d (%d replicates)\n', ...
                scenario.name, spikeFrac, nPerGroup, nReplicates);
        end
    end
end

metadata = metadata(1:row, :);
metadataPath = fullfile(outputDir, metadataCsv);
writetable(metadata, metadataPath);

fprintf('\nDone. Wrote %d count tables to %s\n', row, outputDir);
fprintf('Metadata CSV: %s\n', metadataPath);

function keepIdx = selectTaxaToKeep(counts, taxaKeepFraction)
    nTaxa = size(counts, 2);
    keepM = max(1, round(taxaKeepFraction * nTaxa));
    nnzPerTaxon = sum(counts > 0, 1);
    colSums = sum(counts, 1);
    idx = 1:nTaxa;
    sortMat = [-nnzPerTaxon(:), -colSums(:), idx(:)];
    [~, ord] = sortrows(sortMat, [1 2 3]);
    keepIdx = sort(ord(1:keepM));
end

function nRows = estimateMetadataRows(scenarios, spikeFractions, sampleSizesPerGroup, nReplicates)
    nRows = 0;
    for ii = 1:numel(scenarios)
        nSpikeSettings = numel(spikeFractions);
        nRows = nRows + nSpikeSettings * numel(sampleSizesPerGroup) * nReplicates;
    end
end

function groupLabels = makeGroupLabels(nGroups, nPerGroup)
    groupLabels = zeros(nGroups * nPerGroup, 1, 'uint8');
    for g = 1:nGroups
        startIdx = (g - 1) * nPerGroup + 1;
        endIdx = g * nPerGroup;
        groupLabels(startIdx:endIdx) = g;
    end
end

function baselineContam = buildBaselineContamination(counts, nTaxa, contaminationFraction)
    contamSource = counts(randi(size(counts, 1), 1), :);
    baselineContam = contamSource .* contaminationFraction;
    if numel(baselineContam) ~= nTaxa
        baselineContam = reshape(baselineContam, 1, nTaxa);
    end
end

function eligibleCols = findEligibleTaxa(countsSub, groupLabelsCurr, kPosRequired)
    uniqueGroups = unique(groupLabelsCurr)';
    ok = true(1, size(countsSub, 2));
    for g = uniqueGroups
        rows = groupLabelsCurr == g;
        ok = ok & (sum(countsSub(rows, :) > 0, 1) >= kPosRequired);
    end
    eligibleCols = find(ok);
end

function spikeInfo = initialiseSpikeInfo()
    spikeInfo.indices = {[], [], []};
    spikeInfo.foldChanges = {[], [], []};
    spikeInfo.betas = {[], [], []};
    spikeInfo.alphaEff = [];
end

function spikeInfo = assignScenarioSpikes(scenario, eligibleCols, nSpiked, spikeFoldSetSigned, allowOverlapSpikes, x, alphaEff, groupLabelsCurr)
    spikeInfo = initialiseSpikeInfo();
    spikeInfo.alphaEff = alphaEff;

    if isempty(eligibleCols)
        return
    end

    targetGroups = scenario.groupSpikeTargets;

    if scenario.usesContinuous
        nSelected = min(nSpiked, numel(eligibleCols));
        selected = sort(eligibleCols(randperm(numel(eligibleCols), nSelected)));
        foldChanges = spikeFoldSetSigned(randi(numel(spikeFoldSetSigned), nSelected, 1));
        betas = sign(foldChanges) .* log(abs(foldChanges));
        xNumeric = str2double(x);

        for k = 1:nSelected
            spikeInfo.alphaEff(:, selected(k)) = spikeInfo.alphaEff(:, selected(k)) .* exp(betas(k) .* xNumeric);
        end

        storeGroup = scenario.groupSpikeTargets(1);
        spikeInfo.indices{storeGroup} = selected;
        spikeInfo.foldChanges{storeGroup} = foldChanges;
        spikeInfo.betas{storeGroup} = betas;
        return
    end

    availableCols = eligibleCols;
    for ii = 1:numel(targetGroups)
        groupId = targetGroups(ii);
        nSelected = min(nSpiked, numel(availableCols));
        if nSelected == 0
            continue
        end

        picked = sort(availableCols(randperm(numel(availableCols), nSelected)));
        foldChanges = spikeFoldSetSigned(randi(numel(spikeFoldSetSigned), nSelected, 1));
        betas = sign(foldChanges) .* log(abs(foldChanges));
        groupMask = groupLabelsCurr == groupId;
        multipliers = reshape(exp(betas), 1, []);
        spikeInfo.alphaEff(groupMask, picked) = spikeInfo.alphaEff(groupMask, picked) .* repmat(multipliers, sum(groupMask), 1);

        spikeInfo.indices{groupId} = picked;
        spikeInfo.foldChanges{groupId} = foldChanges;
        spikeInfo.betas{groupId} = betas;

        if ~allowOverlapSpikes
            availableCols = setdiff(availableCols, picked);
        end
    end

end

function txt = summariseGroupCounts(groupLabelsCurr)
    u = unique(groupLabelsCurr)';
    parts = strings(1, numel(u));
    for i = 1:numel(u)
        parts(i) = sprintf('%d:%d', u(i), sum(groupLabelsCurr == u(i)));
    end
    txt = join(parts, ';');
end

function txt = serialiseNumeric(values)
    if isempty(values)
        txt = "";
        return
    end
    txt = join(compose('%.6g', values), ';');
end

function txt = serialiseStrings(values)
    if isempty(values)
        txt = "";
        return
    end
    txt = join(string(values), ';');
end

function txt = scenarioNotes(scenario)
    switch scenario.name
        case "binary_covariate"
            txt = "Binary covariate with spike-ins in group 2 only. Group 1 remains the reference with sample bias, taxa bias, and contamination bias.";
        case "continuous_covariate"
            txt = "Continuous Gaussian covariate with mean 0 and no group distinction. Spike effect is applied as exp(beta*x_i) while sample bias, taxa bias, and contamination bias remain active.";
        case "three_group_spikein"
            txt = "Three-group design with spike-ins only in groups 2 and 3. Group 1 is null but still includes sample bias, taxa bias, and contamination bias.";
        otherwise
            txt = "";
    end
end
```
