## Supplementary Code for "Bias-mitigated microbiome inference refines coronary artery disease signature": parse_index_and_beta_hpc.m

```
function [idx, beta] = parse_index_and_beta_hpc(metaRow, groupId)
    idxField = "spike_taxa_indices_group" + groupId;
    betaField = "spike_beta_group" + groupId;
    idx = parse_list_field_hpc(metaRow, idxField);
    beta = parse_list_field_hpc(metaRow, betaField);
    n = min(numel(idx), numel(beta));
    idx = idx(1:n);
    beta = beta(1:n);
end
```
