## Supplementary Code for "Bias-mitigated microbiome inference refines coronary artery disease signature": parse_list_field_hpc.m

```
function values = parse_list_field_hpc(metaRow, fieldName)
    if ~ismember(fieldName, string(metaRow.Properties.VariableNames))
        values = [];
        return
    end
    raw = string(metaRow.(fieldName));
    if strlength(raw) == 0
        values = [];
        return
    end
    parts = split(raw, ';');
    values = str2double(parts(:)');
    values = values(~isnan(values));
end
```
