## Supplementary Code for "Bias-mitigated microbiome inference refines coronary artery disease signature": parse_metadata_vector_hpc.m

```
function values = parse_metadata_vector_hpc(metadata, fieldName)
    if istable(metadata)
        values = parse_string_field_hpc(metadata.(fieldName));
    else
        values = parse_string_field_hpc(metadata.(char(fieldName)));
    end
end
```
