## Supplementary Code for "Bias-mitigated microbiome inference refines coronary artery disease signature": parse_scalar_field_hpc.m

```
function value = parse_scalar_field_hpc(metaRow, fieldName)
    if ~ismember(fieldName, string(metaRow.Properties.VariableNames))
        value = NaN;
        return
    end
    value = str2double(string(metaRow.(fieldName)));
    if isnan(value)
        value = NaN;
    end
end
```
