## Supplementary Code for "Bias-mitigated microbiome inference refines coronary artery disease signature": parse_string_field_hpc.m

```
function values = parse_string_field_hpc(raw)
    raw = string(raw);
    if strlength(raw) == 0
        values = [];
        return
    end
    parts = split(raw, ';');
    values = str2double(parts(:));
    values = values(~isnan(values));
end
```
