## Supplementary Code for "Bias-mitigated microbiome inference refines coronary artery disease signature": process_dataset_hpc.m

```
function datasetResults = process_dataset_hpc(metaRow, rowIdx, alpha, maxBoot, B_override, useParallel, prescribed_size, prescribed_contam, tmpDir)
    datasetId = string(metaRow.dataset_id);
    fpath = string(metaRow.output_csv);

    if ~isfile(fpath)
        error("Missing count table for %s: %s", datasetId, fpath);
    end

    opts = detectImportOptions(fpath, ...
        'Delimiter', ',', ...
        'ReadVariableNames', true, ...
        'VariableNamesLine', 1, ...
        'DataLine', 2, ...
        'VariableNamingRule', 'preserve', ...
        'TextType', 'string');
    importNames = string(opts.VariableNames);
    metaCols = intersect(["synthetic_sample_id", "source_sample_id", "group", "continuous_covariate"], importNames);
    if ~isempty(metaCols)
        opts = setvartype(opts, cellstr(metaCols), 'string');
    end
    T = readtable(fpath, opts);
    [Y, groupVec, contVec] = extract_design_inputs_hpc(T, datasetId, string(metaRow.scenario_name));
    d = size(Y, 2);

    comparisons = build_comparisons_hpc(metaRow, Y, groupVec, contVec, d);
    if isempty(comparisons)
        error("No valid comparisons constructed for %s", datasetId);
    end

    datasetResults = table('Size', [numel(comparisons) 20], ...
        'VariableTypes', {'string','string','string','double','double','string','double','double','double','double','double','double','double','double','double','double','double','double','double','double'}, ...
        'VariableNames', {'dataset_id','scenario_name','comparison_name','reference_group','target_group','output_csv','spikeFraction','sampleSizePerGroup','rep','nGroups','nSamples','nTaxa','Bused','nTrue','nDiscoveries','TP','Sensitivity','FDR','bootstrapSeconds','percentileSeconds'});

    for c = 1:numel(comparisons)
        cmp = comparisons(c);

        if isempty(cmp.Y) || size(cmp.Y,1) == 0
            error("Empty comparison for %s: %s", datasetId, cmp.name);
        end

        x2 = cmp.X(:,2);
        if all(x2 == 0) || all(x2 == 1)
            error("Degenerate comparison for %s: %s", datasetId, cmp.name);
        end


        metaCmp = subset_metadata_for_rows_hpc(metaRow, cmp.rowIdx);

        out = run_bootstrap_calls_orthog_hpc( ...
            cmp.Y, cmp.X, alpha, maxBoot, B_override, rowIdx + 1000 * c, ...
            useParallel, metaCmp, prescribed_size, prescribed_contam, tmpDir);

        calls = out.calls(:)';
        Gtrue = cmp.Gtrue(:)';

        TP = nnz(calls & Gtrue);
        FP = nnz(calls & ~Gtrue);
        FN = nnz(~calls & Gtrue);
        nDisc = TP + FP;
        sens = TP / max(TP + FN, 1);
        fdr = FP / max(nDisc, 1);

        datasetResults.dataset_id(c) = datasetId;
        datasetResults.scenario_name(c) = string(metaRow.scenario_name);
        datasetResults.comparison_name(c) = cmp.name;
        datasetResults.reference_group(c) = cmp.referenceGroup;
        datasetResults.target_group(c) = cmp.targetGroup;
        datasetResults.output_csv(c) = fpath;
        datasetResults.spikeFraction(c) = parse_scalar_field_hpc(metaRow, "spike_fraction");
        datasetResults.sampleSizePerGroup(c) = parse_scalar_field_hpc(metaRow, "sample_size_per_group");
        datasetResults.rep(c) = parse_scalar_field_hpc(metaRow, "replicate");
        datasetResults.nGroups(c) = parse_scalar_field_hpc(metaRow, "n_groups");
        datasetResults.nSamples(c) = size(cmp.Y, 1);
        datasetResults.nTaxa(c) = d;
        datasetResults.Bused(c) = out.B;
        datasetResults.nTrue(c) = nnz(Gtrue);
        datasetResults.nDiscoveries(c) = nDisc;
        datasetResults.TP(c) = TP;
        datasetResults.Sensitivity(c) = sens;
        datasetResults.FDR(c) = fdr;
        datasetResults.bootstrapSeconds(c) = out.bootstrapSeconds;
        datasetResults.percentileSeconds(c) = out.percentileSeconds;
    end
end
```
