## Supplementary Code for "Bias-mitigated microbiome inference refines coronary artery disease signature": read_metadata_table_hpc.m

function Tmeta = read_metadata_table_hpc(metaFile)
    opts = detectImportOptions(metaFile, ...
        'Delimiter', ',', ...
        'VariableNamingRule', 'preserve', ...
        'TextType', 'string');
    opts = setvartype(opts, opts.VariableNames, 'string');
    Tmeta = readtable(metaFile, opts);
end
