## Supplementary Code for "Bias-mitigated microbiome inference refines coronary artery disease signature": run_bootstrap_calls_orthog_hpc.m

```
function out = run_bootstrap_calls_orthog_hpc(Y, X, alpha, maxBoot, B_override, seed, useParallel, metadata, prescribed_size, prescribed_contam, tempDir)
    if nargin >= 6 && ~isempty(seed), rng(seed); end
    if nargin < 11 || isempty(tempDir), tempDir = tempdir; end

    [~, d] = size(Y);
    if ~isempty(B_override)
        B = B_override;
    else
        alphaAdj = alpha / d;
        B = ceil(200 / alphaAdj);
        if isfinite(maxBoot)
            B = min(B, maxBoot);
        end
    end

    x2 = X(:,2);
    validX2 = x2(~isnan(x2));
    isBinary = ~isempty(validX2) && all(ismember(unique(validX2), [0 1]));

    N = size(X,1);
    if isBinary
        g = 1 + (x2 > 0);
        rowsByGroup = make_rows_by_group_hpc(g);
    else
        rowsByGroup = {};
    end

    fit0 = als_fit_orthog_hpc(Y, X, metadata, prescribed_size, prescribed_contam);
    [p1, dfit] = size(fit0.Ghat);
    chunkSize = min(B, 100);
    alphaAdj = alpha / d;
    pctLo = 100 * (alphaAdj / 2);
    pctHi = 100 * (1 - alphaAdj / 2);

    tmpFile = fullfile(tempDir, sprintf('bootstrap_tmp_%d_%d.mat', seed, randi(1e9)));
    cleanupObj = onCleanup(@() delete_if_exists_hpc(tmpFile));
    mf = matfile(tmpFile, 'Writable', true);
    mf.GhatBoot = zeros(p1, dfit, B);

    tBootstrap = tic;
    if useParallel && chunkSize > 1
        for chunkStart = 1:chunkSize:B
            chunkEnd = min(chunkStart + chunkSize - 1, B);
            nChunk = chunkEnd - chunkStart + 1;
            chunkBoot = zeros(p1, dfit, nChunk);

            parfor jj = 1:nChunk
                b = chunkStart + jj - 1;
                idx = draw_bootstrap_idx_hpc(N, rowsByGroup, isBinary, seed, b);
                fitb = als_fit_orthog_hpc(Y(idx,:), X(idx,:), subset_metadata_for_bootstrap_hpc(metadata, idx), prescribed_size, prescribed_contam);
                chunkBoot(:,:,jj) = fitb.Ghat;
            end
            mf.GhatBoot(:,:,chunkStart:chunkEnd) = chunkBoot;
        end
    else
        for b = 1:B
            idx = draw_bootstrap_idx_hpc(N, rowsByGroup, isBinary, seed, b);
            fitb = als_fit_orthog_hpc(Y(idx,:), X(idx,:), subset_metadata_for_bootstrap_hpc(metadata, idx), prescribed_size, prescribed_contam);
            mf.GhatBoot(:,:,b) = fitb.Ghat;
        end
    end
    bootstrapSeconds = toc(tBootstrap);

    lo = zeros(p1, dfit);
    hi = zeros(p1, dfit);
    tPercentile = tic;
    for jj = 1:dfit
        taxonBoot = mf.GhatBoot(:, jj, :);
        lo(:, jj) = prctile(taxonBoot, pctLo, 3);
        hi(:, jj) = prctile(taxonBoot, pctHi, 3);
    end
    percentileSeconds = toc(tPercentile);
    sig = (lo > 0) | (hi < 0);

    fprintf('  bootstrap loop: %.2fs | percentile loop: %.2fs\n', bootstrapSeconds, percentileSeconds);

    out = struct();
    out.calls = logical(sig(1,:));
    out.B = B;
    out.lo = lo;
    out.hi = hi;
    out.bootstrapSeconds = bootstrapSeconds;
    out.percentileSeconds = percentileSeconds;
end
```
