## Supplementary Code for "Bias-mitigated microbiome inference refines coronary artery disease signature": run_bootstrap_worker_hpc.m

```
clear
clc
close all
rng(1)

projectRoot = pwd;

alpha = 0.05;
maxBoot = inf;
B_override = [];
useParallel = true;
numWorkers = 16;
prescribed_size = 0;
prescribed_contam = 0;

ALS_pair_boot_biased_metadata_orthog_hpc(projectRoot, alpha, maxBoot, B_override, useParallel, numWorkers, prescribed_size, prescribed_contam);
```
