## Supplementary Code for "Bias-mitigated microbiome inference refines coronary artery disease signature": run_eval.R

```
#!/usr/bin/env Rscript

# Resolve script path whether run via Rscript (CLI) or sourced interactively in RStudio
.cmd_file <- grep("^--file=", commandArgs(FALSE), value = TRUE)
if (length(.cmd_file) > 0) {
  script_path <- normalizePath(sub("^--file=", "", .cmd_file[1]), winslash = "/")
} else if (requireNamespace("rstudioapi", quietly = TRUE) && rstudioapi::isAvailable()) {
  script_path <- normalizePath(rstudioapi::getSourceEditorContext()$path, winslash = "/")
} else {
  stop("Cannot determine script path. Either run via 'Rscript run_eval.R' or open this file in RStudio and use Source.")
}
rm(.cmd_file)
project_dir <- dirname(script_path)
source(file.path(dirname(project_dir), "_shared", "simulation_eval_common.R"))

run_project_jobs(project_dir, list(
  list(id = "multimethod", mode = "multimethod", out_subdir = "results",
       methods = c("ancombc2", "maaslin3", "aldex3", "locom", "linda", "wilcoxon"),
       aldex_gamma = 0.5)
))
```
