## Supplementary Code for "Bias-mitigated microbiome inference refines coronary artery disease signature": subset_metadata_for_bootstrap_hpc.m

```
function metadataOut = subset_metadata_for_bootstrap_hpc(metadataIn, idx)
    metadataOut = metadataIn;

    if istable(metadataIn)
        varNames = string(metadataIn.Properties.VariableNames);
    else
        varNames = string(fieldnames(metadataIn));
    end

    if ismember("sample_bias_factors", varNames)
        raw = get_metadata_field_hpc(metadataIn, "sample_bias_factors");
        values = parse_serialised_numeric_hpc(raw);
        if ~isempty(values)
            values = values(idx);
            metadataOut.sample_bias_factors = join(compose('%.6g', values), ';');
        end
    end
end
```
