## Supplementary Code for "Bias-mitigated microbiome inference refines coronary artery disease signature": subset_metadata_for_rows_hpc.m

```
function metaOut = subset_metadata_for_rows_hpc(metaRow, rowIdx)
    metaOut = metaRow;

    if ismember("sample_bias_factors", string(metaRow.Properties.VariableNames))
        sb = parse_list_field_hpc(metaRow, "sample_bias_factors");
        if ~isempty(sb)
            sb = sb(rowIdx);
            metaOut.sample_bias_factors = join(compose('%.6g', sb), ';');
        end
    end
end
```
