## Supplementary Code for "Bias-mitigated microbiome inference refines coronary artery disease signature": README.txt

README

These folders correspond to benchmarking tests performed in our paper. To reproduce results:

1) Open a folder and run 'generate_count_tables.m'.
2) Once complete, run 'run_eval.R' for comparator methods and run 'run_bootstrap_worker_hpc.m' for BootDA adjusting 'useParallel' to 'true' or 'false' as desired to run parallelised bootstrapping and if set 'true' adjust 'numWorkers' to the desired number of CPU cores. Note that BootDA makes use of many bootstrap evaluations and can take time to produce an output.
3) Compile results produced in the corresponding results sub-folders. The compiled set of results for all methods for a given benchmarking scenario that we computed may be found in 'results.xlsx'.
