## Supplementary Information for "Bias-mitigated microbiome inference refines coronary artery disease signature"

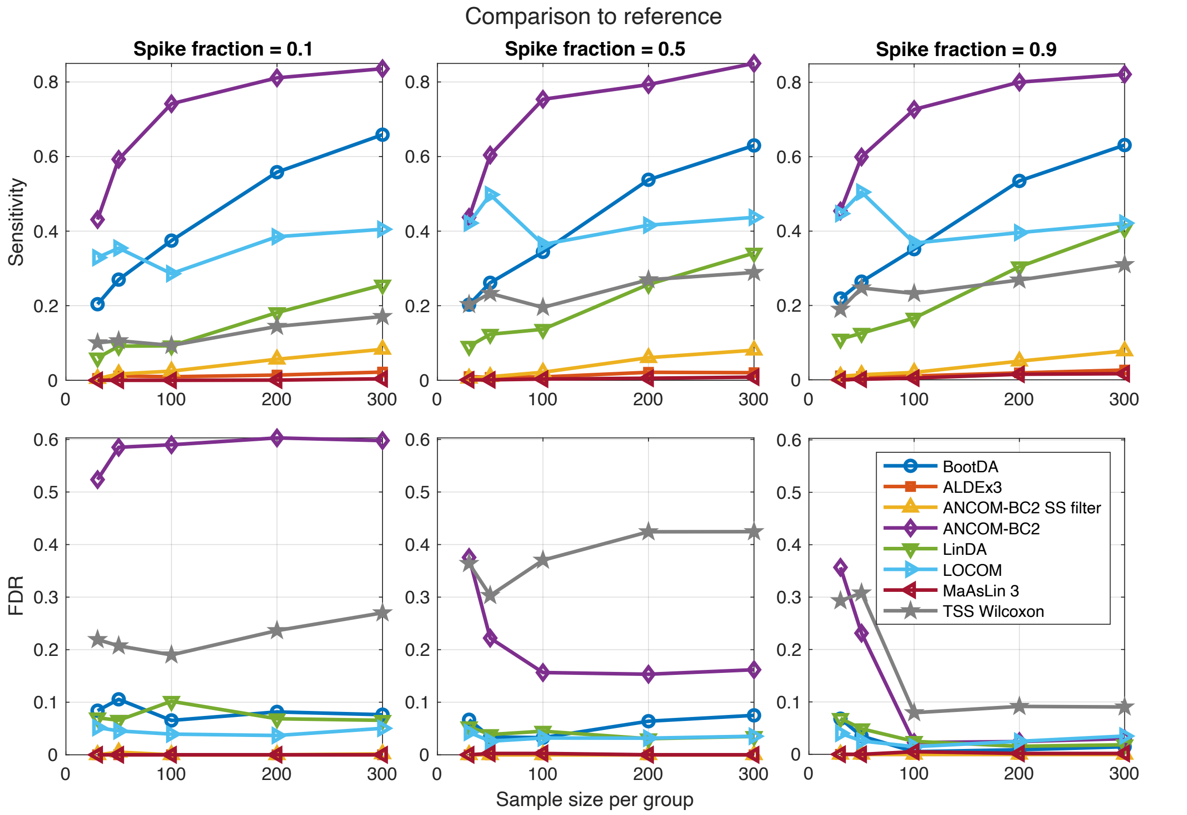


Figure S1. Benchmarking DA methods in simulated Dunnett-style 3-group comparison to reference settings. Mean sensitivity FDR as a function of sample size per group, for BootDA, ALDEx3, ANCOM-BC2 (standard and pseudocount-sensitivity filtered), LinDA, LOCOM, and MaAsLin 3, and Wilcoxon rank-sum testing on relative abundances (TSS Wilcoxon). Means are taken from results over 100 semi-parametric generated count tables derived from the QMP dataset, for spike-in fractions of 0.1, 0.5, 0.9 of taxa.

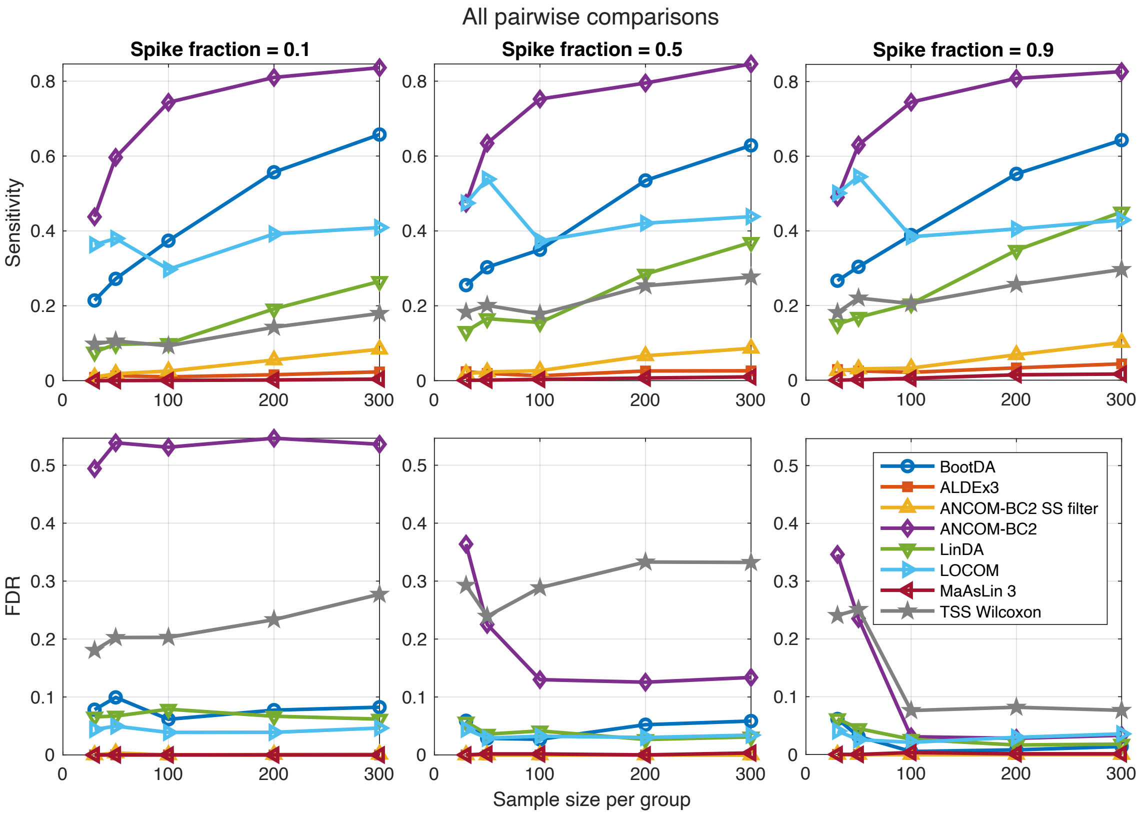


Figure S2. Benchmarking DA methods in simulated 3-group all pairwise comparisons settings. Mean sensitivity FDR as a function of sample size per group, for BootDA, ALDEx3, ANCOM-BC2 (standard and pseudocount-sensitivity filtered), LinDA, LOCOM, and MaAsLin 3, and Wilcoxon rank-sum testing on relative abundances (TSS Wilcoxon). Means are taken from results over 100 semi-parametric generated count tables derived from the QMP dataset, for spike-in fractions of 0.1, 0.5, 0.9 of taxa.

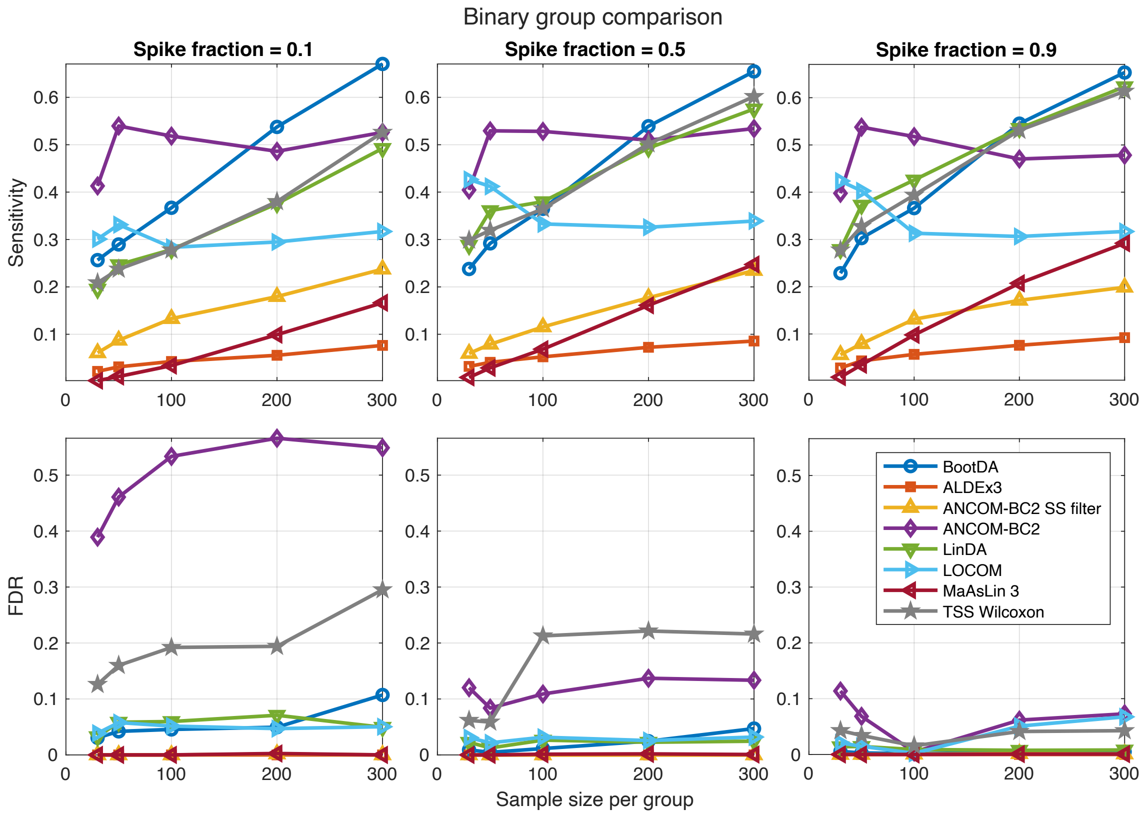


Figure S3. Benchmarking DA methods in binary comparison settings. Mean sensitivity FDR as a function of sample size per group, for BootDA, ALDEx3, ANCOM-BC2 (standard and pseudocount-sensitivity filtered), LinDA, LOCOM, and MaAsLin 3, and Wilcoxon rank-sum testing on relative abundances (TSS Wilcoxon). Means are taken from results over 100 semi-parametric generated count tables derived from the URT dataset, for spike-in fractions of 0.1, 0.5, 0.9 of taxa.
